# Three immunoregulatory signatures define non-productive HIV infection in CD4^+^ T memory stem cells

**DOI:** 10.64898/2026.03.20.713012

**Authors:** Giacomo M. Butta, Bremy Alburquerque, Charlotte Kearns, Yoav Hadas, Max W. VanDyck, Susanna Scaglioni, Noah Peña, Hoi Tong Wong, Elizabeth Levendosky, Charles Gleason, Xiao Lin, Lara Manganaro, Dalila Pinto, Lubbertus C.F. Mulder, Viviana Simon

## Abstract

The persistent HIV reservoir constitutes the main obstacle to curing HIV/AIDS disease. Our understanding of how non-productive HIV infections are established in primary human CD4^+^ T cells during the first round of infection remains, however, incomplete. In this study, we leveraged the HIV reporter virus pMorpheus-V5 to delineate cellular expression patterns that are upregulated in non-productively infected primary CD4^+^ T memory stem cells (T_SCM_). We found that CD4^+^ T_SCM_ harboring non-productive proviruses displayed a distinct transcriptomic signature comprising 118 upregulated genes. This non-productive expression profile was distinct from that of productively infected cells as well as from negative-exposed and mock-infected cells. Among the cellular genes most upregulated in CD4^+^ T cells harboring non-productive proviruses were CCR4-binding migratory chemokines (*CCL22, CCL17*), tryptophan catabolic enzymes (*IDO1, KYNU*), and genes encoding cytoskeletal rearrangement proteins (*BASP1, TNFAIP2*). Intracellular flow cytometry-based analyses confirmed that non-productively infected CD4^+^ T_SCM_ cells were enriched for CCL22 and IDO1 co-expression compared to the other CD4^+^ memory subsets, underscoring a clear CD4^+^ T cell subset specificity for the upregulation of these two immune gene sets associated with non-productive infections. These findings suggest that primary human CD4^+^ T_SCM_ harboring non-productive proviruses display a distinct immunoregulatory phenotype which may facilitate immune evasion and contribute to the persistence of the HIV reservoir.

## Introduction

HIV eradication is challenging because of the persistence of a viral reservoir that evades both immune clearance and antiretroviral therapy (ART)^1–4^. The best characterized persistent reservoir in people with HIV (PWH) is found in long-lived HIV-infected memory CD4^+^ T cells, a highly heterogeneous cell population circulating systemically (e.g., in peripheral blood mononuclear cells, PBMC) or residing in secondary lymphoid tissues^2,5–8^. Estimates suggest that ART-treated PWH with well-suppressed viral replication have less than one in a million circulating CD4^+^ T cells harboring a replication-competent provirus^9,10^. Latently infected cells are heterogeneous cell populations that harbor transcriptionally silent proviruses, which do not produce detectable viral proteins. The lack of viral production is due to post-integration latency resulting from epigenetic silencing of initially productive proviruses as well as from immediate early non-productive infections arising directly after integration. These non-productively infected cells (NP cells) resulting from immediate early latency fail to produce viral proteins despite successful proviral integration^11^.

Most of the HIV latency transcriptomes have been generated from *in vitro* cell culture models in which primary human CD4^+^ T cells are infected with single LTR-dependent reporter viruses and FACS-sorted for productive infection, then cultured for two to eight weeks until epigenetic silencing progressively converts productively infected cells into non-productive ones^12–14^. While these benchmark studies have greatly advanced our understanding of HIV latency, they do not provide information on the properties of primary human CD4^+^ T cells harboring non-productive proviruses established immediately after integration. While reporter viruses are widely used to study the latent reservoir in cell culture model systems, next-generation sequencing (NGS) approaches have provided important insights into the composition of the long-lived persistent HIV reservoirs in PWH, years or even decades after active replication has been stopped^15–18^. However, these approaches do not discriminate between proviruses that were initially productive but subsequently silenced through epigenetic means and proviruses that were never transcriptionally active, which define immediate early latent cells.

To overcome these limitations, dual-reporter HIV constructs have been developed to allow direct and simultaneous *in vitro* detection of cells harboring a non-productive or a productive integrated provirus, based on LTR-dependent and LTR-independent markers^19–22^. HIV-GKO was developed as a single-round virus lacking *nef* and was successfully used to characterize the latent reservoir in both peripheral blood and tonsils^23–25^. HIV pMorpheus-V5 was developed to express all viral genes, including *nef* (with the exception of *env*) and two separate reporter cassettes under the control of either the LTR viral promoter (HSA-mCherry-IRES-Nef cassette) or the cellular PGK promoter (PGK-V5-NGFR cassette)^26^. By including *nef*, the system allows the study of HIV latency under more physiological conditions, as this accessory protein interacts with multiple host factors^27^. Previous studies demonstrated that pMorpheus-V5 efficiently infects primary CD4^+^ T cells, with both productively and non-productively infected cells harboring integrated HIV-DNA^26^. Furthermore, a percentage of non-productive pMorpheus-V5 proviruses remain inducible upon PMA/ionomycin stimulation^28^. For these reasons, the pMorpheus-V5 construct has been widely used to dissect mechanisms of immediate early latency establishment and/or reversal^26,28–31^. To date, however, only a few studies have used these dual reporter viruses to directly analyze the transcriptome of primary human CD4^+^ T cells harboring non-productive proviruses established immediately after integration^21,25^.

CD4^+^ T cell subsets play a central role in establishing long-term immune memory and coordinating rapid recall responses that are essential for controlling the spread of invading pathogens^32^. Cells with a stem cell-like phenotype (T stem cell memory, CD4^+^ T_SCM_) have the longest life span of all CD4^+^ T memory subsets, displaying self-renewal capacity and multipotency^33–35^. These cells continuously regenerate their own pool through homeostatic proliferation and replenish the more differentiated memory CD4^+^ T cell subsets (central memory [T_CM_], transitional memory [T_TM_], and effector memory [T_EM_])^33,34,36^. Prior work from our group and others indicates that CD4^+^ T cell subsets differ in their abilities to support non-productive infections and point to CD4^+^ T_SCM_ as a preferred niche for HIV to persist^37,38,39^. Since the first description of the CD4^+^ T_SCM_ subset, multiple studies have confirmed the pivotal role of this rare subset in HIV infection and persistence^37,40^. Indeed, we and others have reported that, when infected with HIV-GKO dual reporter system^39^, primary CD4^+^ T naïve and T_SCM_ subsets are more prone to immediate early latency (or to harbor a non-productive provirus) when compared to more differentiated subsets. In recent years, *in vivo* data on prolonged ART-treated individuals have gathered evidence of the extreme stability of the CD4^+^ T_SCM_ sub-reservoir, highlighting a critical role in both HIV reservoir establishment and persistence^38,41^.

In the present study, we performed a comprehensive assessment of HIV immediate early latency establishment in primary human CD4^+^ T_SCM_ using the single-round dual reporter HIV pMorpheus-V5 system to identify gene signatures specific for cells harboring non-productive proviruses. Our studies revealed the upregulation of three immunoregulatory pathways, linked to CCR4-binding migratory chemokines, tryptophan catabolic enzymes and cytoskeletal rearrangement, as specific markers for immediate early latency establishment. Furthermore, NP CD4^+^ T_SCM_ cells were specifically enriched for a sub-population co-expressing CCL22 and IDO1.

## Results

### HIV pMorpheus-V5 detects non-productively infected CD4^+^ T cells

Bulk analysis of HIV-infected CD4^+^ T cells can miss virus-induced signatures specific for smaller cell populations, as in the case of CD4^+^ T cells harboring non-productive HIV proviruses. Therefore, we leveraged the HIV dual reporter virus pMorpheus-V5^26^ (Figure 1a) to distinguish between CD4^+^ T cells harboring non-productive proviruses (termed “NP” phenotype cells) and productively infected cells (termed “P” phenotype cells) or negative-exposed bystander cells, which have encountered virus but remain uninfected (“NE” phenotype cells).

**Figure 1.**
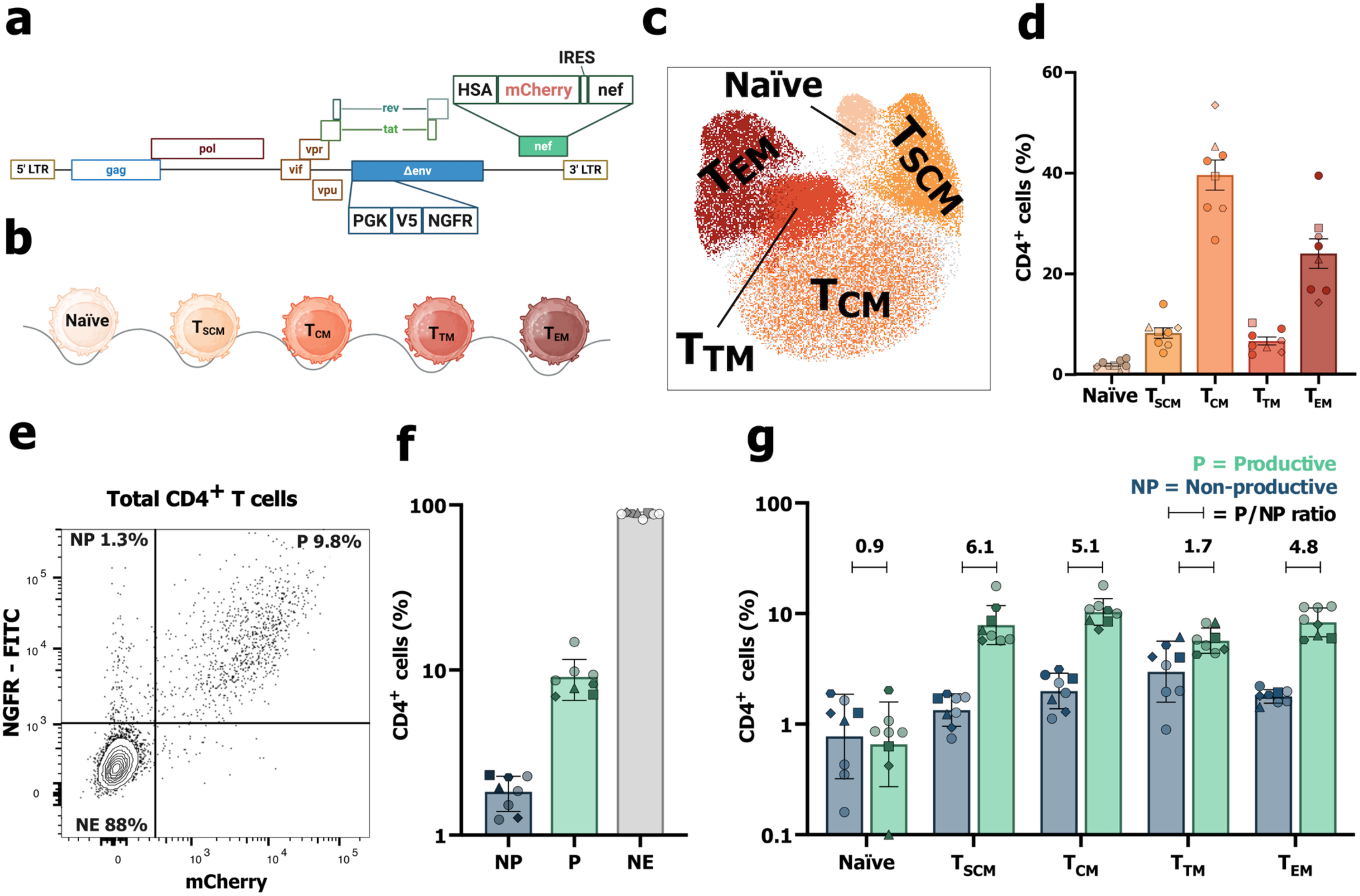
HIV pMorpheus-V5 dual-reporter virus discriminates between non-productive and productive infections in primary CD4^+^ T cell subsets. (a) Representation of HIV pMorpheus-V5 genome, with the LTR-dependent (HSA-mCherry-IRES-Nef) and LTR-independent (PGK-V5-NGFR) cassettes highlighted. (b) Representation of peripheral blood CD4^+^ T naïve and memory subsets (T memory stem cells [T_SCM_], central memory [T_CM_], transitional memory [T_TM_] and effector memory [T_EM_]). (c) Representative Uniform Manifold Approximation and Projection (UMAP) of 211,934 cells overlaying five CD4^+^ T cell subsets (naïve, T_SCM_, T_CM_, T_TM_, T_EM_) within total CD4^+^ T cell population (gated on live cells, CD4^+^ T cell subsets are color-coded), at five days post-infection (dpi). (d) Bar graph showing the frequencies for each CD4^+^ T cell subset within total live CD4^+^ population (n=8), at five dpi. (e) Representative contour plot showing FACS gating strategy to define non-productively infected (NP), productively infected (P) and negative-exposed (NE) CD4^+^ T cells (cells gated on total CD4^+^ cells), defined by NGFR and mCherry expression. (f) Frequencies of the NP, P and NE populations in total CD4^+^ T cells, at the time of FACS analysis (five days post-infection, n=8). (g) Percentages of non-productively and productively infected cells in each CD4^+^ T cell subset considered (n=8) with the ratio of productive over non-productive (P/NP ratio) highlighted for each subset. Each symbol represents a different healthy blood donor, with the four donors used for subsequent CD4^+^ T_SCM_ sorting highlighted with different symbols. Donors used for analysis but not sorted are represented by paler dots. Primary CD4^+^ T cells were stimulated with αCD3/CD28 antibodies for 72 hours before infection with HIV pMorpheus-V5 single-round reporter virus for five days before analysis. Data presented as the mean with SD.

To explore the heterogeneity of the non-productive HIV cellular landscape during the first round of replication, we coupled the HIV pMorpheus-V5 system with flow cytometry to discriminate between CD4^+^ T naïve cells and CD4^+^ memory T cell subsets (T_SCM_, T_CM_, T_TM_ and T_EM_; Figure 1b). Briefly, CD3/CD28-activated peripheral blood CD4^+^ T cells were infected by spinoculation^42^ with the single-round HIV pMorpheus-V5 reporter virus pseudotyped with a dual-tropic subtype B env (n=8 different healthy anonymous blood donors). Five days post-spinoculation, we observed that, among the different subsets, CD4^+^ T_CM_ were the most abundant (Figure 1c and 1d, 39.6% of total live CD4^+^ T cells on average, range: 26.7 – 53.5%), while naïve CD4^+^ T cells were the least abundant (Figure 1d, 2% on average, range: 0.7 – 3.3%). CD4^+^ T_SCM_ made up 8.2% (range: 4.3% – 14%) of all CD4^+^ T lymphocytes (Figure 1d). The proportion of infected cells (NGFR^+^mCherry^-^ and NGFR^+^mCherry^+^) was significantly lower in the CD4^+^ T naïve subset compared to CD4^+^ memory subsets (p<0.0001 compared to T_CM,_ Supplementary Figure 1b). For CD4^+^ T_SCM_ this difference was less prominent (p=0.033, compared to T_CM_), while no significant differences were observed between CD4^+^ T_CM_, T_TM_ and T_EM_ (Supplementary Figure 1b).

Next, we quantified the fraction of each CD4^+^ T cell subset with respect to pMorpheus-V5 infection phenotypes (non-productive [NP, NGFR^+^mCherry^-^], productive [P, NGFR^+^mCherry^+^] cells, Figure 1e). Single-cycle infection of total CD4^+^ T cells resulted in an average of 1.8% NP (range: 1.2 - 2.3%), 9.1% P (range 6.9 - 14.8%) and 88.0% of NE (Figure 1f, range: 81.7 - 90.1%). Infection of naïve CD4^+^ T cells was comparably low for both NP and P (Figure 1g, 1% of NP vs. 0.8% of P, ratio P/NP 0.9) while CD4^+^ T_TM_ had the highest frequency of NP (Figure 1g, 3.4% of NP vs. 5.9% of P, ratio P/NP 1.7). CD4^+^ T_CM_ and CD4^+^ T_EM_ displayed very similar P/NP ratios (5.1 and 4.8, respectively). The CD4^+^ T_SCM_ subset showed 1.4% of NP (range 0.7 - 1.9%) and 7.1% of P (Figure 1g, range 5.7 - 17.7%, P/NP ratio 6.1). Notably, infection of naïve CD4^+^ T cells was the only instance in which NP cells were comparable to P cells (Figure 1g, ratio P/NP 0.9).

Taken together, these data suggest that immediate early non-productive infections occur in CD4^+^ T cell subsets with different frequencies. NP cells are present in each of the CD4^+^ T cell subsets analyzed with limited donor variability, allowing for robust downstream interrogations of these long-lived CD4^+^ T cells.

### CD4^+^ T_SCM_ cells harboring non-productive proviruses express a distinct gene expression signature

We next tested the hypothesis that the transcriptomic profiles of CD4^+^ T_SCM_ cells vary according to pMorpheus-V5 infection outcomes. Briefly, infected CD4^+^ T_SCM_ from four healthy donors (donors 1, 2, 3 and 4) were sorted based on infection phenotypes (Figure 2a and Supplementary Figure 1a). Percentages of NP, P and NE cells in both total CD4^+^ T cells and in CD4^+^ T_SCM_ cells were comparable to previous experiments (total CD4^+^: NP range 1.3 - 2.2%, P range 6.9 - 8.2%. CD4^+^ T_SCM_: NP range 0.9 – 1.9%, P range 5.7 – 11.3% and NE range: 88.9 - 90.0%, respectively, Supplementary Figure 1c, 1d and 1e). Mock-infected cells were sorted as negative controls (termed “mock”). Cellular RNA and DNA was extracted from sorted cells. Digital PCR was performed on DNA to assess proviral intactness (Figure 3). Poly(A) enrichment was performed on the RNA and paired-end NGS libraries were prepared. 13 – 37 million RNA-seq reads per sample were analyzed to identify expression patterns specifically associated with non-productive infection.

**Figure 2.**
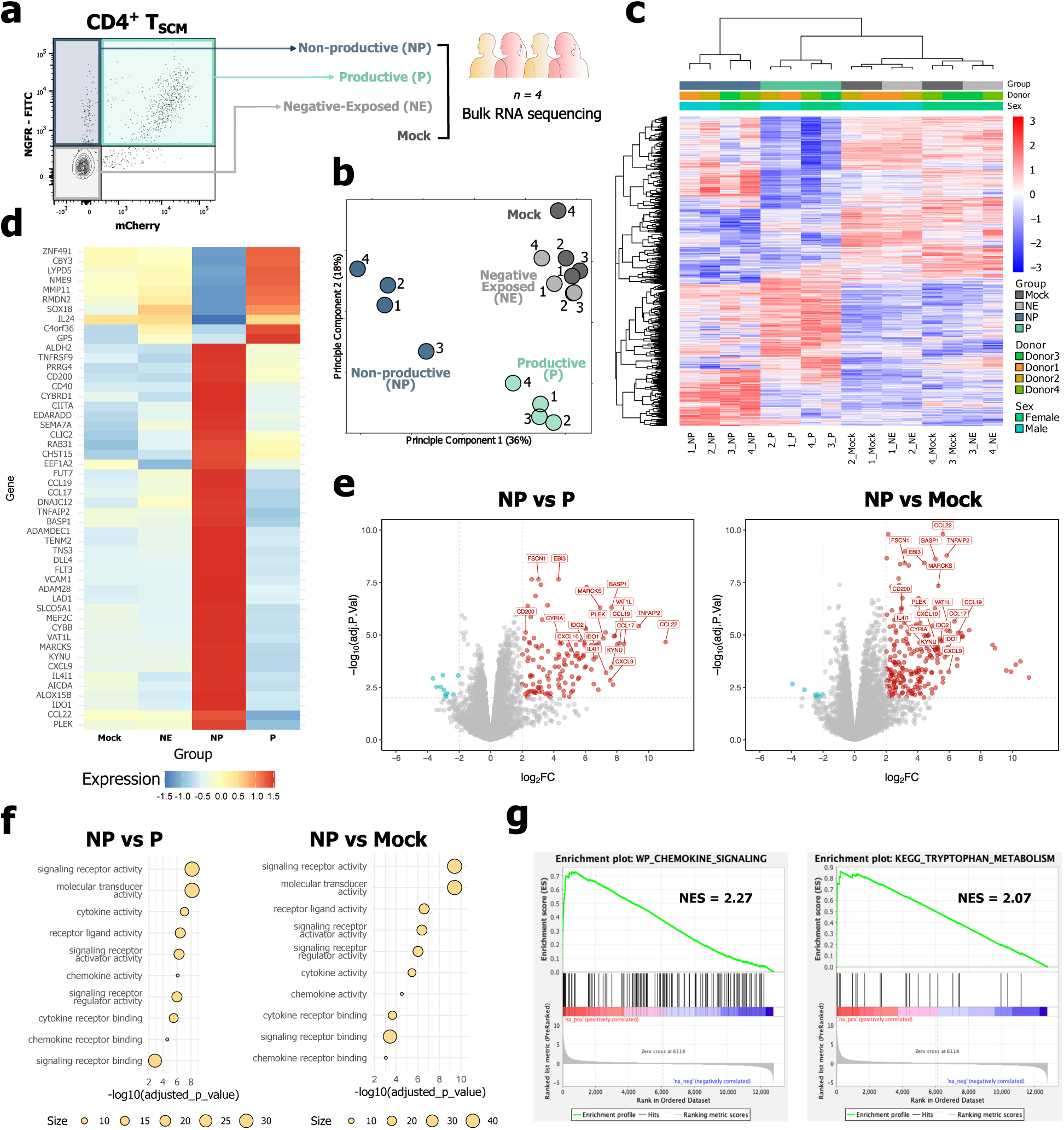
Non-productively infected CD4^+^ T_SCM_ transcriptome exhibits chemokine production, tryptophan catabolism activity and cytoskeletal rearrangement signature. (a) Experimental strategy. Freshly isolated CD4^+^ T cells from peripheral blood were spinoculated with the pMorpheus-V5 dual reporter virus. Five days post-infection, cells were stained with a panel of antibodies to discriminate between CD4^+^ T cell subsets, and CD4^+^ T_SCM_ were sorted based on the infection outcome (NP, P, NE or mock). RNA and DNA were extracted from lysed cells and bulk RNAseq was performed (n=4). (b) Principal component analysis (PCA) of gene expression pattern in sorted populations (NP, P, NE, mock) of four donors (donor 1-2-3-4). (c) Unsupervised clustering heatmap with dendrograms from bulk RNA sequencing (RNAseq) showing the expression levels (Z-score) of differentially expressed genes across sorted populations and donors. (d) Heatmap of selected differentially expressed genes (Z-score is shown, row normalized considering only the displayed genes) across all sorted CD4^+^ T_SCM_ populations. (e) Volcano plots representing the relative expression of all genes, with highlighted genes encoding for CCR4-binding chemokines, tryptophan catabolism enzymes, cytoskeletal rearrangement and other relevant genes in NP vs P (left) and NP vs mock (right) comparisons. 131 genes upregulated in NP vs P, 185 upregulated in NP vs mock, (log_2_FoldChange (FC) >= 2, FDR <= 0.01). (f) Gene ontology (GO) molecular function (MF) of significantly upregulated genes in NP CD4^+^ T_SCM_, compared to P (left) and mock (right). (g) Pre-ranked gene set enrichment analysis (GSEA) for genes differentially expressed between NP and P sorted populations, using a molecular signature of chemokine signaling (left) and tryptophan metabolism (right). Normalized enrichment score (NES) is shown.

**Figure 3.**
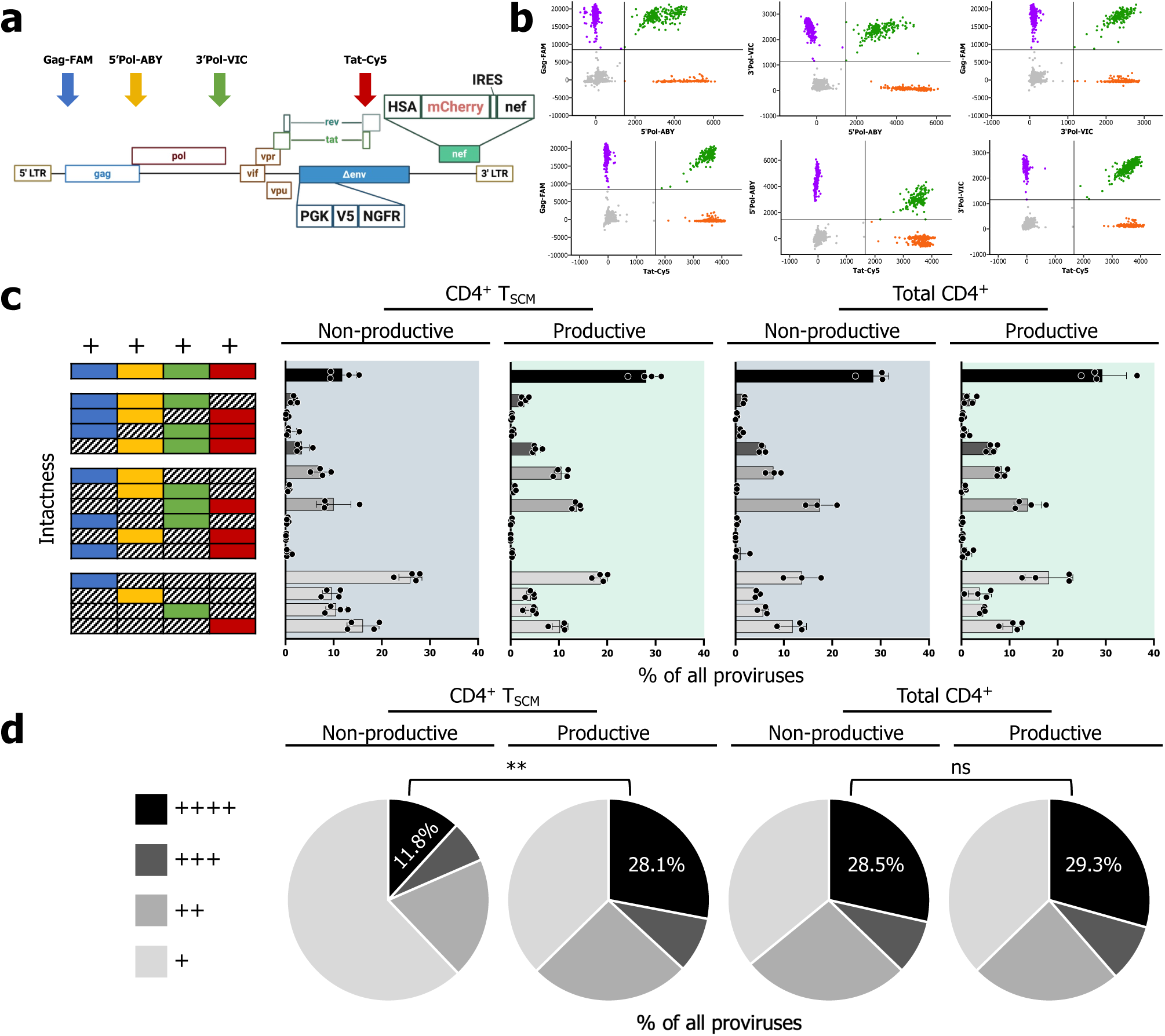
dPCR-Proviral Intactness Assay (dPCR-PIA) demonstrates intact HIV proviruses within sorted non-productively and productively infected CD4^+^ T cells. (a) Schematic of pMorpheus-V5 reporter virus with arrows denoting the location of 4 probes, Gag-FAM, 5’Pol-ABY, 3’Pol-VIC, and Tat-Cy5. (b) Representative data from dPCR with single and double positive wells of each probe pair. (c) dPCR-PIA was run on DNA isolated from infected and sorted for non-productive and productive proviruses, from either sorted CD4^+^ T_SCM_ or total CD4^+^ T cells. Schematic to the left depicts the permutations of probe combinations while the bar graphs to the right show the % of positive wells that correspond to each combination averaged across four donors (or three donors for total CD4^+^ NP). (d) Pie charts summarize the percentages of proviruses which are called as intact (++++) or positive for any 3, 2, or 1 probe. The brackets above denote the significance of paired t-test comparing the fraction of intact in NP vs P across paired donors (ns, Not Significant, *, p<0.05, **, p<0.01).

Principal component analyses (PCA) of the four cell subsets originating from the four different donors (Figure 2b) indicated that infection phenotypes strongly influenced the cellular transcriptome, while inter-donor variability was less influential. Negative-exposed cells (NE) clustered close to the mock control cells (mock), indicating that exposure to the single-round HIV pMorpheus-V5 did not substantially alter gene expression. To identify specific patterns across the groups, we performed differential gene expression analysis using limma (DGE, false-discovery rate corrected p-value (FDR) <= 0.01). Consistent with the PCA analysis, unsupervised clustering analysis confirmed that each group (NP, P, NE and mock) clustered separately from the other, except for NE and mock populations, which overlapped (Figure 2c). The comparison of differentially expressed genes (DE genes) between NE and mock populations did not reveal any significant changes (Supplementary Figure 2a). Importantly, NP populations were transcriptionally different from P, NE and mock, displaying a cluster of 118 commonly upregulated genes (log_2_FC >= 2 and FDR <= 0.01, Figure 2d and Supplementary Figure 2c). In NP CD4^+^ T_SCM_ cells, CCR4-binding chemokines (*CCL22* and *CCL17*) transcripts exhibited the highest fold increase (log_2_FC = 11.1 and 8.4, compared to P, log_2_FC = 5.6 and 6.3 compared to mock, Figure 2e), alongside other genes involved in T cell migration and cytoskeletal rearrangement (*CCL19*, *CXCL9*, *CXCL10*, *EBI3*, *TNFAIP2*, *FSCN1*, *MARCKS*, *PLEK* and *BASP1*, Figure 2d and 2e). *CCL22* and *CCL17* transcripts were expressed almost exclusively in NP populations (Figure 2d and Supplementary Figure 2e). *TNFAIP2* (or M-Sec) showed the highest log_2_FC among actin-binding genes (average: 9.4) compared to P cells, followed by another actin-binding protein (*BASP1*, average: log_2_FC = 7.7, Figure 2e left). Both genes were exclusively expressed in NP populations (Figure 2d and Supplementary Figure 2e). In addition, transcripts encoding tryptophan catabolic enzymes were consistently and exclusively upregulated in NP (*IDO1*, *IDO2*, *KYNU*, *IL4I1*, Figure 2d, 2e, and Supplementary Figure 2e). Productively infected CD4^+^ T_SCM_ cells compared with other populations showed a transcriptomic difference (selected genes in Supplementary Figure 2d), and a stringent set of DE genes when compared to mock populations only (74 upregulated genes, eight downregulated genes, |log_2_FC| >= 2 and FDR <= 0.01, Supplementary Figure 2b). To assess the presence of virus-specific DE genes, we analyzed overlapping upregulated genes found exclusively in P and NP populations compared to NE and mock (Supplementary Figure 2f). Only six genes were shared by NP and P: *CD22*, *SPATA18*, *LMNA*, *SERPINE1*, *ARHGAP31* and *TSKU*.

To complement the DGE analysis, we performed functional enrichment analyses of the NP CD4^+^ T_SCM_ population using gProfiler2 with the annotations of Gene Ontology (GO) and Kyoto Encyclopedia of Genes and Genomes (KEGG) (Figure 2f and Supplementary Figure 2g). Molecular function (MF) was strongly enriched in categories associated with secretion (signaling receptor activity, cytokine activity, chemokine activity, adj. p-value range 7.1×10^-9^ - 7.9×10^-7^ compared to P and adj. p-value range 4×10^-10^ - 3×10^-5^ compared to mock, Figure 2f), while the gene set enrichment analysis (GSEA) showed significant enrichment scores for the tryptophan metabolism set and the chemokine signaling set (Normalized Enrichment Score (NES) = 2.07 and 2.27, respectively, Figure 2g). Biological processes (BP) of NP CD4^+^ T_SCM_ cell populations were enriched in cell adhesion and cell migration (adj. p-value range 1.5×10^-10^ - 8.8×10^-9^ compared to P and adj. p-value range 2.96×10^-11^ - 3.96×10^-11^ compared to mock, Supplementary Figure 2g).

Together, CD4^+^ T_SCM_ harboring non-productive proviruses are characterized by a transcriptomic landscape that is markedly different from that of productively infected, negative-exposed or mock-infected cells. This NP signature is characterized by upregulation of three gene sets: CCR4-binding chemokines, tryptophan catabolic enzymes, and actin-binding proteins. Of note, none of these pathways are active in P, NE or mock-infected control CD4^+^ T_SCM_.

### Validation of the three immunoregulatory gene sets in total CD4^+^ T cells

We next investigated whether the transcriptomic signature identified in NP CD4^+^ T_SCM_ cells was also detectable in a broader population, such as total CD4^+^ T cells exhibiting the NP infection phenotype. To this end, we sorted CD4^+^ T cells from five additional healthy donors separating NP, P and NE populations (donors 5, 6, 7, 8 and 9, Supplementary Figure 3a). To ensure the highest purity of the NP cell population, cells were stained for the HSA membrane protein, which is included in the LTR-dependent cassette in addition to mCherry (Figure 1a). Therefore, NP CD4^+^ T cells were HSA^-^ (in addition to NGFR^+^mCherry^-^), while P CD4^+^ T cells were HSA^+^ (in addition to NGFR^+^mCherry^+^, Supplementary Figure 3a). The overall infection rates of these experiments were consistent with previous experiments (NP range: 0.8 – 1.4%, P range: 7 – 12.3%, NE: 83.1 – 89.7%, Supplementary Figure 3b). The expression levels of the six genes most upregulated in NP CD4^+^ T_SCM_ were verified by reverse transcription quantitative PCR (RT-qPCR) using RNA extracted from sorted CD4^+^ T cells (Supplementary Figure 4a, 4b and 4c). CCR4-binding chemokines (*CCL22* and *CCL17*), one tryptophan catabolic enzyme (*KYNU*) and cytoskeletal rearrangement genes (*BASP1* and *TNFAIP2*) were significantly upregulated in NP CD4^+^ T cells compared to P CD4^+^ T cells (p=0.02, p=0.005, p=0.01, p=0.008, p=0.02, respectively). *IDO1* was also upregulated in NP cells but due to inter-donor variation the differences did not reach statistical significance (p=0.09 compared to P). When compared to mock-infected cells, expression levels of the six signature genes were always significantly upregulated in NP CD4^+^ T cells (except for IDO1), while the comparison to NE reached significance in 50% of the analyses (*CCL22*, *KYNU*, *BASP1*, Supplementary Figure 4a, 4b and 4c). These results demonstrate that the lead candidate genes defining the immunoregulatory NP CD4^+^ T_SCM_ signature were consistently upregulated in total CD4^+^ T cells harboring non-productive proviruses.

### Absence of non-productive gene signatures in HIV-negative publicly available single-cell datasets

We further examined whether the top upregulated genes in NP cells were co-expressed at the single-cell level in peripheral blood CD4^+^ T cells or other immune populations from healthy individuals, alongside ten additional upregulated genes implicated in T cell migration, cytoskeletal remodeling, and tryptophan catabolism. For this analysis, we utilized two publicly available single-cell RNA-seq (scRNA-seq) datasets: polarized peripheral blood CD4^+^ T cells (CD3/CD28-activated naïve and memory CD4^+^ T cells in the presence or absence of different combinations of cytokines)^43^ and the Human Immune Health Atlas^44^ (Supplementary Figure 5). Within the peripheral blood CD4^+^ T cell dataset, simultaneous expression of the six top-ranked genes (*CCL22, CCL17, IDO1, KYNU, TNFAIP2*, and *BASP1*) was restricted to a single Th17 cell out of the total 43,112 cells. Furthermore, only 37 cells co-expressed three or more of these genes (Supplementary Figure 5c); these were predominantly identified as Th17 (n=16) and iTreg (n=17) populations (Supplementary Figure 5b). These findings were further confirmed in the Human Immune Health Atlas (Supplementary Figure 5d-f). Among more than 1.8 million cells spanning 29 immune cell types, no individual cells were found to express either the six top genes or the expanded 16-gene panel simultaneously. Conventional type 1 dendritic cells (cDC1s) exhibited the strongest enrichment for the NP gene set; however, only a small fraction of cells (21 of 943; 2%) expressed incomplete and variable combinations of the six top genes and 14 of the 16 extended genes (Supplementary Figure 5f). Overall, these results show the absence of a subset of circulating immune cells known to co-express the transcriptomic signature characterizing NP CD4^+^ T_SCM_ cells.

### NP and P CD4^+^ T cells carry near-intact proviruses

We leveraged our pMorpheus-V5 specific multiplex digital PCR proviral intactness assay (dPCR-PIA) to evaluate the intactness of proviruses after a single round of infection in NP and P CD4^+^ T cell populations, in both sorted CD4^+^ T_SCM_ and total CD4^+^ T cells. Briefly, our dPCR-PIA allows for partition of single integrated proviruses into 20,480 individual microwells (AbsoluteQ, Thermofisher), where quantitative PCR reactions occur, with four sets of primers and probes corresponding to distinct regions of the HIV genome (e.g., Gag, 5’Pol, 3’Pol, Tat, Figure 3a). Microwells positive for all four probes are scored as putatively intact proviruses (Figure 3b and 3c), in line with the data interpretation implemented in the Intact Proviral DNA Assay, an established protocol that relies on the detection of two proviral regions to call intactness. Briefly, we extracted genomic DNA simultaneously with RNA from sorted P and NP CD4^+^ T_SCM_ as well as P and NP total CD4^+^ T cells (four different donors each, see Supplemental Table 2). We observed that 11.4% (range: 9.3 - 15.3%) of the CD4^+^ T_SCM_ with a non-productive phenotype harbored intact proviruses compared to 28.1% (range: 27.7 - 31.2%) of the productively infected CD4^+^ T_SCM_ cells (p=0.007; paired T test, Figure 3c and d). In contrast, in total CD4^+^ T cells, the fraction of intact proviruses in NP and P infected cells was comparable (NP: 28.5% (range: 24.8 - 30.4%) intact proviruses vs P: 29.3% (range: 24.9 - 36.5%) in the P infected cells (p=0.580; paired T test, Figure 3c and d).

### CCL22 or L-Tryptophan treatment does not perturb latency establishment

Chemokines and substrates of the tryptophan metabolism pathway highlighted in the DGE analysis of NP CD4^+^ T_SCM_ cells could act in an autocrine manner to modulate latency establishment. To investigate this aspect, we treated activated primary CD4^+^ T cells with CCL22 or L-Tryptophan (Trp) prior to infection with pMorpheus-V5 and then quantified the levels of NP and P populations (Supplementary Figure 6a). In mock-treated total CD4^+^ T cells, the ratio of P/NP cells was 8.8, with CCL22 or Trp-treated cells showing comparable P/NP ratios (8.8 and 8.7, respectively, Supplementary Figure 6b). At the T cell subset level (Supplementary Figure 6c to g), P/NP ratios were all comparable (ratios of 2, 1.5 and 1.8 for CCL22, Trp, mock treatment in naïve cells; ratios of 8, 9.1 and 8.1 for CCL22, Trp, mock treatment in T_SCM_ cells; ratios of 8.1, 7.7 and 7.4 for T_CM_ cells; ratios of 2.5, 3.3, 3.4 for T_TM_ and ratios of 5.4, 5.4 and 6.2 for the T_EM_). These findings indicate that CCL22 and tryptophan do not influence the proportion of non-productively and productively infected cells in primary human CD4^+^ T cells.

### CCL22^+^ IDO1^+^ cells are enriched in NP CD4^+^ T_SCM_ cells

To obtain single-cell resolution protein expression data, we developed a 13-color flow-cytometry panel to measure intracellular expression of CCL22 and IDO1 in five CD4^+^ T cell subsets. We found a significant enrichment of CCL22^+^IDO1^+^ cells (Figure 4a and 4b, single positive and double positive populations shown) within the NP CD4^+^ T_SCM_ cells subset compared to P, NE and mock cells (p=0.04, p=0.03, p=0.04 respectively, Figure 4a and 4d). On average, 8.3% of NP CD4^+^ T_SCM_ cells co-expressed CCL22 and IDO1 (range 6.5 - 10.9%), compared to 0.3% of P cells (range 0 - 0.6%), 0.5% NE (range 0.02% – 0.99%) and 0.1% mock cells (Figure 4d, range 0.07 - 0.2%). Of note, no significant differences were found between the NP populations and P, NE or mock populations within the other CD4^+^ T subsets analyzed (Figure 4c, e, f, g). In total CD4^+^ T cells, NP cells displayed a significant increase in the levels of CCL22^+^IDO1^+^ cells (average of 1.6%, range 1.3 - 1.9%), compared to P (average of 0.2%, range 0.2 - 0.3%, p=0.02), NE (average of 0.05%, range 0.02 - 0.06%, p=0.01) and mock (Figure 4h, average of 0.04%, range 0.02 - 0.07%, p=0.01). When we compared only NP populations across the different CD4^+^ T cell subsets, CCL22^+^IDO1^+^ cells were more prevalent in CD4^+^ T_SCM_ (Figure 4i, p=0.02 compared to naïve, ns compared to other subsets). Notably, the expression of these two intracellular proteins in CD4^+^ T_SCM_ was highly correlated, with the vast majority co-expressing CCL22^+^ and IDO1^+^ (Figure 4a, single IDO1^+^CCL22^-^: 0%, IDO1^-^CCL22^+^: 1.56%, IDO1^+^CCL22^+^: 10.9%, IDO1^-^CCL22^-^: 87.5%).

**Figure 4.**
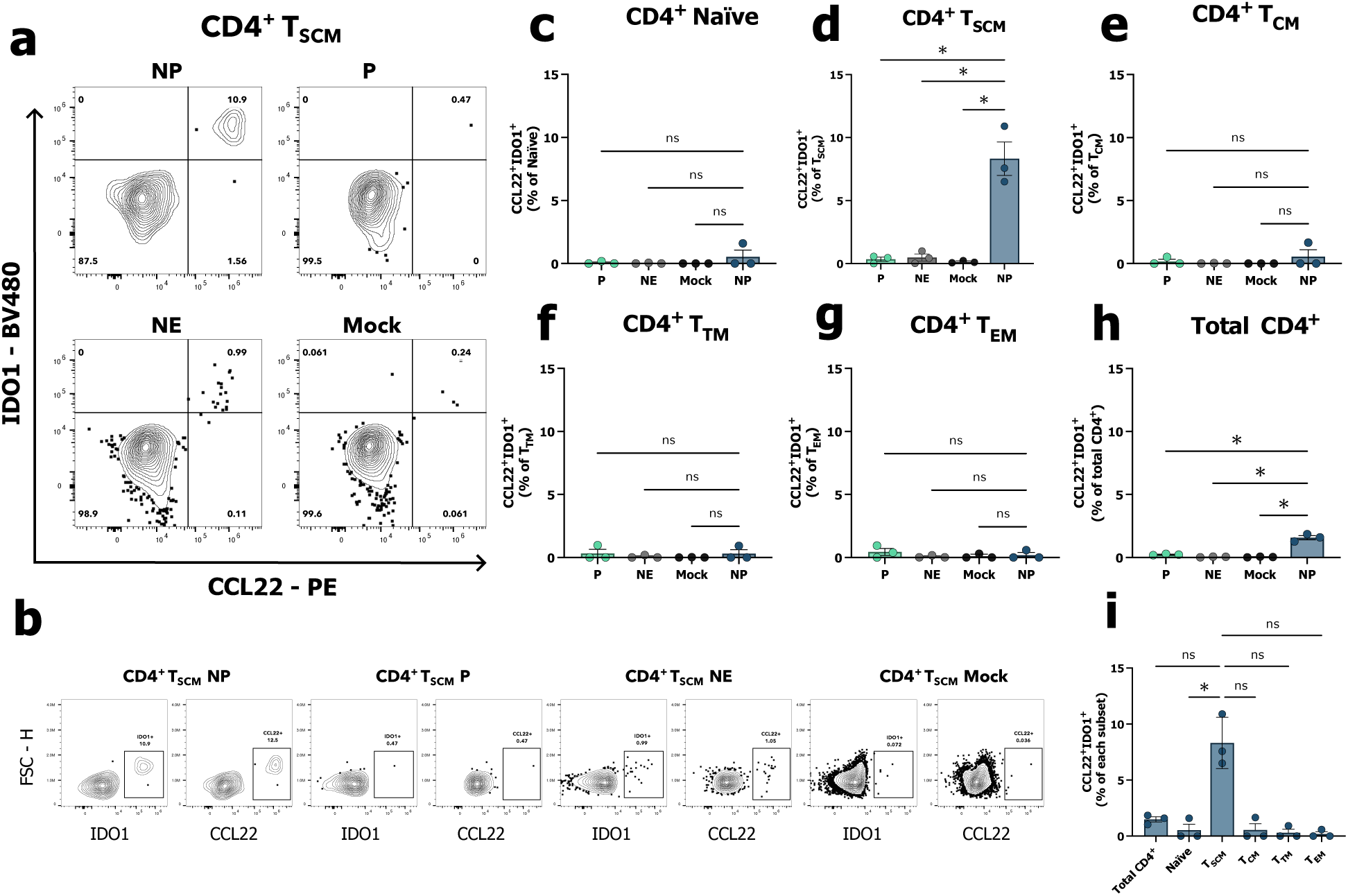
CCL22^+^IDO1^+^ cells are enriched in NP CD4^+^ T_SCM_ cells. (a) Representative contour plots gated on CD4^+^ T_SCM_ population showing the percentage of CCL22^+^IDO1^+^ cells in NP, P, NE and mock populations. (b) Representative contour plots gated on CD4^+^ T_SCM_ population showing the percentage of single positive populations for either IDO1 or CCL22, in CD4^+^ T_SCM_ NP, P, NE and Mock. (c through h) Bar graphs displaying the percentages of CCL22^+^IDO1^+^ cells in every subset considered, in naïve (c), T_SCM_ (d), T_CM_ (e), T_TM_ (f), T_EM_ (g) and total CD4^+^ T cells (h, n=3). (i) Bar graph showing the percentage of CCL22^+^IDO1^+^ in NP population across the CD4^+^ T cell subsets considered (n=3). Significance was calculated using Repeated Measures one-way ANOVA with multiple comparisons (Dunnett’s multiple comparisons test, between NP, P, NE and Mock). *, P < 0.05: **, P< 0.01, ***, P < 0.001, ****, P < 0.0001, ns, not significant. Data presented as the mean with SD.

These data demonstrate that the non-productively infected cells in total CD4^+^ T cells are significantly enriched for cells co-expressing CCL22 and IDO1. Notably, this pro-tolerogenic phenotype is most prominently represented within the T_SCM_ subset.

## Discussion

Our study shows that non-productive HIV infections are associated with the activation of specific cellular pathways during the first round of replication. Transcriptomic analyses detected three major signatures characterizing CD4^+^ T_SCM_ bearing non-productive proviruses: CCR4-binding chemokines (*CCL22* and *CCL17*) alongside other chemokines (*CCL19*, *CXCL9*, *CXCL10*, *EBI3*), tryptophan catabolic enzymes (*IDO1*, *IDO2*, *KYNU*, *IL4I1*) and factors involved in cytoskeletal rearrangement (*TNFAIP2, BASP1, FSCN1*, *MARCKS*, *PLEK*, *CYRIA*). These gene sets and proteins are fundamentally involved in immunoregulatory processes, facilitating regulatory T cells (Tregs) interactions with target cells and driving differentiation of regulatory phenotypes^45–47^. When we analyzed transcript levels across conditions, we confirmed that this upregulation is specific to NP cells. Since only a limited number of studies have examined the CD4^+^ T cell transcriptomic landscape of immediate early latency using dual-reporter HIV vectors so far^21,25^, our transcriptomic datasets represent a valuable resource for future characterizations of non-productive infections.

*IDO1*, the gene that encodes indoleamine 2,3-dioxygenase 1, was one of the most significantly upregulated genes in NP CD4^+^ T_SCM_ cells. IDO1 is the rate-limiting enzyme that catalyzes the conversion of tryptophan to kynurenine and has a pivotal role in both immunoregulation and tryptophan metabolism^48^. Alongside *IDO1,* NP CD4^+^ T_SCM_ showed the upregulation of other enzymes (*IL4I1*, *KYNU*, *IDO2*) that convert tryptophan into other metabolites following two different pathways, the I3P and the Kynurenine pathways. Cells expressing IDO1 suppress innate and adaptive immunity to promote tolerance by catabolizing tryptophan to other indole compounds^49^. In the HIV context, elevated IDO1 activity is associated with multiple phenotypes, such as the loss of Th17/Treg balance and the inhibition of HIV replication via tryptophan depletion^50,51^. At the molecular level, IDO1 and IL4I1-generated metabolites act as endogenous ligands of the aryl hydrocarbon receptor (AhR), activating the AhR pathway that leads to FOXP3 expression and Treg differentiation^52,53^. *CCL22* and *CCL17* were also significantly upregulated in NP CD4^+^ T_SCM_. By binding to CCR4, both CCL22 and CCL17 facilitate the recruitment of Tregs to sites of inflammation^54^. The presence of Tregs can inhibit local immune responses, thereby potentially fostering conditions conducive to the establishment and persistence of HIV latency^55^. CCL22 is also secreted by B cells in lymph nodes to attract T cells, foster interactions, and induce germinal center formation^56^. Moreover, CCL22 is secreted by dendritic cells (DCs) in the lymph nodes to attract Tregs, increasing cell-to-cell contacts and initiating an immunosuppressive cascade^46^.

Non-productive infection of CD4^+^ T_SCM_ was associated with the expression of genes involved in cytoskeletal dynamics and cell motility. Actin-associated proteins such as TNFAIP2, FSCN1, MARCKS, PLEK and BASP1 have been shown to cover multiple roles in many cell types^57–61^, converging on the induction of immunological synapses, filopodia and membrane protrusions. More specifically, TNFAIP2 is known to facilitate cell-to-cell HIV transmission via tunneling nanotubes formation^62^, while MARCKS and BASP1 exert their functions in cell protrusions through interactions with the PIP_2_ phospholipid^59,63^. In addition, some evidence points to FSCN1 as the cytoskeletal component underlying Treg cell-mediated suppression in a contact-dependent manner^47^. The specific upregulation of these genes in NP cells indicates a phenotype characterized by cytoskeletal rearrangements. Such changes are absent in productively infected CD4^+^ T_SCM_ and shed light on possible pathways of immune evasion mediated by cell motility and intercellular adhesion.

The three gene sets enriched in NP cells could be markers for a subset of CD4^+^ T cells prone to harbor non-productive proviruses, or a transcriptomic rewiring induced by non-productive proviruses. By analyzing two publicly available scRNAseq datasets, we demonstrated that only a negligible portion of peripheral blood single-cell co-expresses *CCL22*, *CCL17*, *IDO1*, *KYNU*, *BASP1* and *TNFAIP2*, suggesting that a distinct transcriptional program predisposing cells to harbor a non-productive provirus is unlikely.

There is currently little data on intactness after a single round of HIV replication. Bruner et al. estimated 59% intact proviruses after two days of infection in CD4^+^ T cells infected with HIV Ba-L and treated with enfuvirtide^64^. We confirmed the presence of intact pMorpheus-V5 proviruses in our sorted samples using a pMorpheus-specific multiplex dPCR-based proviral intactness assay (dPCR-PIA). The dPCR-PIA builds upon the well-established IPDA protocol for measuring HIV proviral intactness^65^. In our experiments, the relative levels of proviral intactness measured in NP and P cells were low, between 11-30% of all proviruses. These numbers could be an underestimation, as sheared DNA may decrease the number of wells that are positive for all four probes. However, reports indicate that people with primary HIV infection or people with HIV initiating ART early harbor less than 10% intact proviruses within a few weeks of infection^64,66^. These data together suggest that the size of the intact HIV reservoir is quite small from the onset of the infection.

When we treated CD4^+^ T cells with exogenous CCL22 or tryptophan, the proportion of either NP or P cells did not change, indicating that this immunoregulatory profile is a downstream consequence of harboring a non-productive provirus rather than a driver of it. While various chemokines have been linked to HIV latency^67,68^, direct evidence connecting CCL22 and CCL17 to HIV latent infection remains scarce. Research on chemokines with similar physiologic functions, such as CCL19 and CCL21, indicates that chemokine-induced signaling may promote HIV nuclear localization and integration in resting CD4^+^ T cells^67,68^. On the other hand, CCL22 and CCL17 are upregulated in the plasma of ART-treated PWH and secreted during HIV primary immune response^69,70^. As tryptophan alone was not able to modify the establishment of NP cells, it is still unclear whether IDO1-mediated tryptophan depletion, the production of its metabolites, or their signaling through the Kyn-AhR axis has an impact on immediate early HIV latency in CD4^+^ T cells^71,72^.

We successfully confirmed the NP transcriptomic signature at the protein level using flow cytometry: a distinct fraction of NP CD4^+^ T_SCM_ co-expressed CCL22 and IDO1. While this immunoregulatory signature was detectable in total CD4^+^ T cells via both flow cytometry and qPCR in sorted populations, it was disproportionately enriched in the CD4^+^ T_SCM_ subset when compared to the other subsets. It is attractive to speculate that this subset-specific concentration of CCL22 and IDO1 may establish an immunosuppressive niche, potentially limiting the elimination of non-productively infected cells and facilitating the persistence of latent reservoirs. In this scenario, NP CD4^+^ T_SCM_ could recruit Tregs by secreting CCL22 and then leverage the immunosuppressive capabilities of these Tregs to maintain the NP phenotype at the single-cell level. At the same time, IDO1 activity in T_SCM_ harboring a non-productive provirus could generate immunosuppressive kynurenines and sustain this environment through multiple mechanisms.

In conclusion, we identified three gene sets unique to non-productively infected CD4^+^ T_SCM_ cells that share immunosuppressive features absent in either productive infections or exposed uninfected bystander cells. At the protein level, co-expression of CCL22 and IDO1 was highly enriched in non-productively infected CD4^+^ T_SCM_. We suggest that non-productive proviruses have an active role in rewiring infected cells in a specific manner that results in upregulation of a set of genes that skew the infected cell and/or its surrounding environment towards an immunosuppressive phenotype, which facilitates immune evasion and promotes persistence. Further studies will elucidate whether an immunosuppressive cellular environment plays a role in HIV persistence, possibly akin to the immunosuppressive tumor microenvironment that inhibits anti-cancer immune responses and facilitates cancer cell survival.

## Materials and methods

### Primary CD4^+^ T cell isolation and culture

Buffy coats were obtained from anonymous healthy donors from the New York Blood Bank (New York Blood Center Component Laboratory). Peripheral blood mononuclear cells (PBMCs) were purified through density gradient centrifugation (Lymphoprep, Stemcell Technologies # 07851) and CD4^+^ T cells were isolated through negative selection (Miltenyi, CD4^+^ T cell Isolation Kit human # 130-096-533) according to the manufacturer’s instructions. Purified CD4^+^ T cells were kept in culture in RPMI medium 1640 (Gibco, # 21870-076) supplemented with 100 U/mL Penicillin-Streptomycin (Corning, # 30-002-CI), 2 mM L-glutamine (Corning, # 25-005-CI), MEM non-essential Amino Acids (Corning, # 25-025-CI), 10 mM HEPES Buffer (Corning, # 25-060-CI) and 10% (vol/vol) heat-inactivated FBS (Gemini Bio, # 100-106). For CD4^+^ T cell stimulation, cells were resuspended in RPMI and IL-2 (Miltenyi, 30 IU/mL # 130-097-744) or activated by adding magnetic beads coated with αCD3/CD28 monoclonal antibodies (Gibco, DynaBeads Human T-activator CD3/CD28 # 11131D, 1:10 ratio beads:cells) and IL-2 30 U/mL for 72 hours. Beads were then separated by placing cells on a magnetic stand, and transferred cells were kept in culture in RPMI and IL-2 (30 IU/mL) for 5 days before FACS analysis or sorting. Cells were maintained at 37°C in a 5% CO_2_ humidified incubator.

### Virus production

HEK 293T cells (ATCC#: CRL-3216) were maintained in DMEM media (Corning, Dulbecco’s Modification of Eagle’s Medium 1X, # 10-013-CV) with the addition of 10% (vol/vol) heat-inactivated FBS, 100 U/mL Penicillin-Streptomycin, MEM non-essential amino acids, and 2 mM L-glutamine. Cells were cultured at 37°C in a 5% CO_2_ humidified incubator. HIV pMorpheus-V5 viral stocks were generated by PEI (polyethylenimine) transfection of HEK 293T cells with 20 μg of pLAI2-Δenv-V5-NGFR-mCherry plasmid (pMorpheus-V5, described in Kim et al.^26^) and 5 μg of X4-R5 dual-tropic envelope pSV-III-92-HT 593.1 plasmid (BEI resources, catalog number # ARP-3077). Sixteen hours post-transfection, the cell culture media was replaced with fresh DMEM (10% FBS), and supernatants were collected 40 hours and 64 hours post-transfection, centrifuged at 500 x g for 5 minutes and stored clarified at 4°C until concentration. Clarified supernatants were filtered through a 0.45 μm PES filter and concentrated approximately 150-fold using LentiX concentrator (ClonTech, Takara Bio # 631232), following the manufacturer’s instructions. Concentrated virus was resuspended in PBS and treated with DNase I (NEB, 100 U/mL, # M0303S) and Benzonase nuclease (Millipore Sigma, 25 U/mL, # 70746-3) for 1 hour at 37°C following the manufacturer’s instructions, then aliquoted and stored at −80°C. p24 viral antigen was quantified using HIV-1 p24 ELISA kit (Xpress Bio, # XB-1000) and optical density was collected using a microtitration plate reader (Victor, Perkin Elmer Victor). Infectious virus titer was calculated through serial dilutions of the viral stock on TZM-bl reporter cell line and quantification of β-Galactosidase signal was collected using the microtitration plate reader as previously described^73^.

### Infection of primary human CD4^+^ T cells

Purified CD4^+^ T cells were seeded at 1×10^6^ cells/mL and stimulated for 3 days in RPMI + αCD3/CD28 magnetic beads and IL-2 30 U/mL. αCD3/CD28 beads were removed before proceeding as directed by the product manufacturer. Subsequently, cells were seeded in RPMI with the addition of polybrene 2 μg/mL (Sigma-Aldrich, # TR-1003) in a U-bottom 96-well plate and infected by spinoculation for 2h at 800 x g, at 32°C with 100 ng of p24 per 10^6^ cells (adapted protocol from O’Doherty et al., JVI 2000)^42^. Spinoculation media was removed, and complete RPMI + IL-2 30 U/mL was added to the cells. Culture supernatant was replaced every 2 days by fresh RPMI + IL-2 30 U/mL until flow cytometry analysis.

### Flow cytometry staining

For flow cytometry analysis, 1×10^6^ primary CD4^+^ T cells infected with pMorpheus-V5 reporter virus or mock were stained and analyzed on a FACS Symphony A5 machine (BD Biosciences) or Cytek Aurora (Cytek). Viability dye Near-IR (Invitrogen, Cat # L34976A) was used to discriminate between live and dead cells. Membrane staining was performed by incubating cells at 37°C for 20 minutes with the following antibodies: αCD45RA (BioLegend, # 568712), αCD45RO (BioLegend, # 304206), αCD95 (BioLegend # 305624), αCD27 (BD, # 565116), αCCR7 (BD, # 566602), αCD271 (BioLegend, # 345104). Cells were washed and resuspended in PBS + 2% FBS and then analyzed. For intracellular staining, 1×10^6^ primary CD4^+^ T cells infected with pMorpheus-V5 reporter virus or mock were treated with BD GolgiStop (containing 0.26% Monensin, BD Biosciences, #554724) for 4 hours, at day 5 post-infection. Subsequently, cells were washed and stained for surface markers together with live/dead near-IR dye, at 37°C for 20 minutes: αCD45RA (BD, # 612927), αCD4 BUV395 (BD, # 564724), αCD45RO (BD, # 748367), αCD95 (BioLegend # 305624), αCCR7 (BD, # 566602), αHSA (BioLegend # 101819), αCD271 (BioLegend, # 345104), αCD27 (BD, # 565116). Then, cells were fixed, permeabilized and stained for intracellular chemokines (αCCL22, BD # 565950, αIDO1, BD # 568019) using the Cytofix/Cytoperm kit (BD Biosciences, # 554714). Flow cytometry analysis was performed using FlowJo software (BD Biosciences, v10).

### Cell sorting

For CD4^+^ T_SCM_ cell sorting, between 50 and 100×10^6^ freshly isolated primary CD4^+^ T cells were stimulated with αCD3/CD28 antibodies for 72 hours, infected with HIV pMorpheus-V5 (100 ng of p24 per 10^6^ cells) for five days and then stained for surface antigens (αCD45RA, αCD45RO, αCD95, αCD27, αCCR7, αCD271-NGFR) and live/dead dye (Near IR) for 20 minutes at 37°C (antibodies catalogs previously described). Cells were washed twice in PBS + 2% FBS, filtered through a 50 μm filter and kept on ice until sorting. Stained cells were gated for CD4^+^ T_SCM_ (CD95^+^, CD45RA^+^, CD45RO^-^, CCR7^+^, CD27^+^) and sorted into non-productively infected, productively infected, negative-exposed and mock unexposed. NGFR^+^/mCherry^-^ CD4^+^ T_SCM_ cells were considered non-productively infected, while NGFR^+^/mCherry^+^ CD4^+^ T_SCM_ were considered productively infected. NGFR^-^/mCherry^-^ CD4^+^ T_SCM_ were considered negative-exposed, while CD4^+^ T_SCM_ mock unexposed cells never experienced the virus, and they were sorted based on being negative for live/dead dye (Near IR) and positive for the T_SCM_ markers mentioned above. A 4-way sorting was performed on an S6 Symphony sorter (BD Biosciences) using a 100 μm nozzle. Sorted cells were lysed the same day in RLT plus buffer (part of the Allprep DNA/RNA Micro kit, Qiagen, # 80204) and stored at −80°C until RNA/DNA extraction.

For total CD4^+^ T cell sorting, between 25 and 70×10^6^ of freshly isolated primary CD4^+^ T cells were stimulated with αCD3/CD28 antibodies for 72 hours, infected with HIV pMorpheus-V5 (100 ng of p24 per 10^6^ cells) for 5 days and then stained for surface antigens (αCD4, αHSA [BioLegend, # 101820], αCD271-NGFR) and live/dead dye (Near IR) for 20 minutes at 37°C. NGFR^+^/mCherry^-^/HSA^-^ CD4^+^ T cells were considered non-productively infected, NGFR^+^/mCherry^+^/HSA^+^ CD4^+^ T cells were considered productively infected, while negative-exposed were negative for all three markers (NGFR^-^/mCherry^-^/HSA^-^). Mock unexposed cells were sorted based on being negative for LD NIR stain. Cells were washed twice in PBS + 2% FBS, filtered through a 50 μm filter and kept on ice until sorting, which was carried out in the same way as previously described for CD4^+^ T_SCM_ sorting. Sorted populations (NP, P, NE and mock unexposed) were then lysed right after sorting in RLT plus buffer and stored at −80°C until RNA/DNA extraction. Flow cytometry analysis was performed using FlowJo software (BD Biosciences, v10).

### RNA isolation and library preparation for RNA-seq

Cellular RNA and DNA from FACS-sorted CD4^+^ T_SCM_ or total CD4^+^ T cells were isolated using Allprep DNA/RNA Micro kit (Qiagen, # 80204). RNA was treated on column with DNase I (Qiagen, # 79254) and quantified using Qubit HS RNA assay kit (Invitrogen, # Q32852). RNA quality was determined using Agilent RNA 6000 Nano Kit (Agilent, # 5067-1511) on the Agilent Bioanalyzer 2100 instrument. Samples with RNA integrity number (RIN) > 8 were used for library preparation. RNA-seq libraries were produced using the SMART-Seq mRNA LP (with UMIs) kit (Takara Bio, # 634762), following the manufacturer’s instructions and starting from an input of 10 ng of total extracted RNA per sample. Along with sorted RNA samples, a positive control (intact RNA of known concentration provided by the kit) and a negative control (nuclease-free water, provided by the kit) were included in each reaction. RT-cDNA was amplified through 10 PCR cycles, and amplified cDNA was purified (AMPure XP Beads, Beckman Coulter, # A63882) and run on Agilent 2100 Bioanalyzer using Agilent High Sensitivity DNA Kit (Agilent, # 5067-4626). cDNA input was normalized at 2.5 ng before library prep reactions and fragmented libraries were PCR amplified through 15 PCR cycles. Amplified libraries were singularly purified (AMPure XP Beads, Beckman Coulter), run on Agilent 2100 Bioanalyzer using Agilent High Sensitivity DNA Kit and quantified at Qubit using Qubit dsDNA BR Assay kit (Invitrogen, # Q33265). Individual libraries were normalized at 10 nM and pooled together. Molarity of the pooled library was confirmed through KAPA Library Quantification PCR (KAPA Biosystems, # KK4824). Pooled libraries were sequenced on a NovaSeqX Plus instrument (Illumina) using a paired-end sequencing approach with reads of 150 base pairs in length (PE150 strategy).

### RNA-Seq data processing, quantification, and differential gene expression analysis

Paired-end SMART-seq reads were first processed using the “demux” command from the Cogent NGS Analysis Pipeline (v2.0.1, Takara Bio) to identify barcodes and UMIs and remove UMI-associated sequences from reads. Further read processing and differential expression analysis were performed using a custom pipeline^74,75^. Briefly, reads were trimmed for adapter sequences and low base quality (quality score threshold < 20 at ends) using cutadapt^76^. Quality control metrics calculated using RNA-seQC^77^, PicardTools (“Picard Toolkit.” 2019. Broad Institute, GitHub Repository: https://broadinstitute.github.io/picard/; Broad Institute), and samtools^78^ were aggregated into multiQC^79^ and manually inspected to ensure sufficient library diversity. Genotype concordance checks used varscan^80^. The STAR aligner (v. v2.7.11b)^81^ was used to map processed reads to a combined genome reference containing both the human genome (assembly GRCh38.p13) and the HIV pMorpheus-V5 genome. Human genes were annotated against GENCODE release v41 and gene count summaries were made using featureCounts^82^, and a raw count matrix was generated with genes as the rows and samples as the columns. Genes with expression levels above 0.5 FPKM in at least 10% of samples were retained for further analysis. The weighted trimmed mean of M-values (TMM) method was used to calculate normalization factors on the filtered matrix and the voom^83^ mean-variance transformation was applied to prepare the data for differential expression analysis with limma^84^. The data were fitted to a design matrix that contained all sample groups (i.e., non-productive, productive, negative-exposed, and mock). Donors were treated as a random effect in the linear model, and the sex of the donor was added as a covariate with a fixed effect. Pairwise comparisons were made between all groups, and empirical Bayes adjusted p-values were corrected for multiple testing using the Benjamini-Hochberg (BH) method to identify differentially expressed genes (FDR q<0.01). Additional fold-change thresholds were applied as indicated.

### Gene Ontology, Gene Set Enrichment Analysis and Pathway Enrichment Analysis

Enrichment for Gene Ontology (GO; biological process, molecular function, and cellular component) and Kyoto Encyclopedia of Genes and Genomes (KEGG) pathways was performed using the gProfiler2 R package (v0.2.3). The background gene set was restricted to the genes in our final filtered gene expression matrix. Differentially expressed genes were ranked by log_2-_fold-change and used as an ordered query. Multiple testing correction was performed using the default g:SCS algorithm. Gene set enrichment analysis (GSEA) was performed using the GSEA software (UC San Diego and Broad Institute). The normalized enrichment score (NES) was calculated by the software from the enrichment score (ES) as it follows: NES = actual ES/mean (ESs against all permutations of the dataset).

### Digital PCR-Proviral Intactness Assay

Digital PCR-Proviral Intactness Assay (dPCR-PIA) is a method to approximate proviral intactness with single molecule resolution, similarly to the previously described and accepted intact proviral DNA assay (IPDA)^65^. dPCR-PIA utilizes a digital PCR system where the sample is distributed across 20,480 microwells, and 4 probes in Gag, 5’ Pol, 3’ Pol and Tat, compared to 2 probes in IPDA corresponding to the HIV packaging site and Env (Primer/Probe sequences in Supplemental Table 2). Approximate primer location sourced from Levy et al., 2021^85^, adapted to pMorpheus-V5 sequence. Briefly, genomic DNA was extracted from sorted CD4^+^ T cells (NP, P, E, mock) using the Allprep DNA/RNA Micro kit (Qiagen, # 80204). Paired RNA samples were used in RNAseq and RT-qPCR experiments. DNA was quantified using the Qubit dsDNA BR Assay Kit (Invitrogen, # Q32853). For each sample, 5 ng of genomic DNA was combined with Absolute Q DNA Digital PCR Master Mix (1X final, Thermo Fisher #A52490), four primer pair and probe sets (900nM and 250nM final, respectively) and diluted in PCR-grade water. Samples were loaded into MAP16 Digital PCR plates (Thermo Fisher # A52865) and topped with isolation buffer and loaded into the machine following the manufacturer’s instructions. Samples were run on the Quantstudio Absolute Q dPCR (Thermo Fisher # A52864), with a preheat cycle of 10 minutes at 96°C, and 40-cycles of 5 seconds at 96°C, and 15 seconds at 60°C. Analysis was performed using the Quantstudio Absolute Q Software Version 6.3. Thresholds were set based on each fluorescent probe and kept consistent across samples. Software identifies positive probes in each of the 20,480 microwells per sample. Data were reported as a percentage of positive wells which are single, double, triple, or quadruple positive for any of the four probes.

### Single-cell RNA-Seq expression and enrichment analysis

Two single-cell RNA sequencing (scRNA-seq) datasets of human immune cells were used to assess enrichment of NP infection-associated genes: 1) the scRNA-seq (10X 3’ v2) Cano-Gamez *et al*.^43^ dataset, consisting of naïve and memory T (TM) cells isolated from PBMCs of four healthy adult males (average 56.4 years of age, SD 12.41 years) and stimulated with five different cytokine combinations. 2) the Human Immune Health Atlas^44^ with 1.82 million cells that includes all major human immune cell types isolated from PBMCs of 108 healthy individuals (age range 11-65 years).

The Cano-Gamez dataset^43^ was obtained from Open Targets^86^ (https://www.opentargets.org/projects/effectorness) in MTX format along with cell metadata and Uniform Manifold Approximation and Projection (UMAP) coordinates. These files were converted into Seurat objects and analyzed using Seurat v5^87^. The Human Immune Health Atlas dataset^88^ was obtained in AnnData (.h5ad) format^88^, which included both cell metadata and UMAP coordinates. Initial processing and subsetting were performed with Scanpy^89^, after which the data were converted to Seurat objects using Scanpy and Seurat v5. Subsequent analyses were conducted with Seurat v5 to generate UMAP visualizations, dot plots, and heatmaps.

Enrichment analysis of 16 key genes expressed in NP infected cells was performed using the UCell scoring method^90^ by assigning an enrichment score to each cell based on the Mann-Whitney U statistic, with higher scores indicating stronger enrichment for the gene set under investigation.

### cDNA synthesis and quantitative real-time RT-qPCR

Isolated cellular RNA was reverse-transcribed (RT) to cDNA using iScript cDNA Synthesis kit (Bio-Rad, # 1708890), normalizing RNA input for all samples. Gene expression was measured by quantitative real-time PCR (qPCR) using iQ SYBR Green Supermix (Bio-Rad, # 1708880) by the comparative quantification cycle (Cq) method according to the manufacturer’s instructions on the Azure Cielo qPCR instrument (Azure Biosystems). β-actin (ACTB) was used as a housekeeping gene to normalize the data. The limit of detection was set at Quantification cycle (Cq) = 38. qPCRs were carried out in triplicate. A table of primer sequences used is provided as Supplementary Table 2.

### Treatment of CD4^+^ T cells with CCL22 or L-Tryptophan

Primary CD4^+^ T cells were isolated and TCR activated following the protocol described in the “Primary CD4^+^ T cell isolation and culture” section and spinoculation as described in the “Infection of primary human CD4^+^ T cells” section. Six hours before spinoculation, CCL22 100 ng/mL (MedChemExpress, # HY-P72790) or tryptophan 50 ug/mL (Sigma, # T8941) was added to the culture media. After spinoculation, the infection media was replaced with RPMI including IL-2 30 IU/mL and either CCL22 (100 ng/mL) or tryptophan (50 ug/mL) or no additions. The media was changed every two days. Five days post-infection, cells were stained and analyzed at FACS following the protocol described in the section “Flow cytometry staining” (only live-dead and surface markers).

### Quantification and statistical analysis

Statistical analyses were performed using Prism 10 software (GraphPad, v10.5.0) or Limma package (v 3.58.1). Details of statistical analysis are provided in each figure legend, including sample size and statistical test used. Volcano plots were produced using the R package ggplot2 (v3.5.2).

## Acknowledgements

We thank the members of the Mount Sinai flow cytometry core (Martina Di Verniere, Sabrina La Salvia, Jovvian George Parakkal, Alessandro Marins Dos Santos and Guillermo Adolfo Villegas) for the assistance during sorting experiments. We thank Marcia Slater (Thermo Fisher Scientific) and Yang Lu for their assistance with the dPCR. This work was supported, in part, by NIH/NIDA R61 DA058294 (L.C.F.M.) and NIH/NIAID R01AI179598 (V.S.). Additional support for training was provided to C.K. and N.P. by NIH/NIAID T32AI007647. This work was also supported in part through the Minerva computational and data resources and staff expertise provided by Scientific Computing and Data at the Icahn School of Medicine at Mount Sinai and supported by the Clinical and Translational Science Awards (CTSA) grant UL1TR004419 from the National Center for Advancing Translational Sciences. Research reported in this publication was also supported by the Office of Research Infrastructure of the National Institutes of Health under award number S10OD030463. The content is solely the responsibility of the authors and does not necessarily represent the official views of the National Institutes of Health.

## Author contributions

V.S., L.C.F.M., and D.P. conceived the experiments. G.M.B., S.S., C.K., M.V.D., E.L., H.T.W., L.C.F.M. performed the experiments and the investigation. B.A., X.L., N.P., C.G., D.P. performed the sequencing data analysis. B.A., D.P., G.M.B., N.P. followed the sequencing data curation. V.S., G.M.B., L.C.F.M., D.P., L.M., Y. H. undertook data interpretation. G.M.B., S.S., B.A., H.T.W. curated the methodology. G.M.B., B.A., D.P. performed the data visualization. V.S., L.C.F.M., D.P. secured funding acquisition, expertise and resources. G.M.B., V.S. wrote the manuscript. V.S., L.C.F.M., D.P. reviewed and edited the manuscript. All authors approved the final manuscript version.

## Declaration of interest

The authors declare no competing interests

## Supplemental Appendix

**Supplementary Table 1.** Summary of donors’ demographics and experiments performed. The table provides an overview of metadata of the healthy donors from whom primary CD4^+^ T cells were used for the experiments described in the manuscript.

| Buffy coat code | Donor # | Sex | Age | Data type | Sorted | Sorted Cell type | Data shown in figure |
| --- | --- | --- | --- | --- | --- | --- | --- |
| 43 | 1 | Male | 31 | FACS analysis, RNAseq, dPCR | Yes | CD4<sup>+</sup> T<sub>SCM</sub> | Figure 1, 2, 3, Supp 1, Supp 2 |
| 68 | 2 | Male | 66 | FACS analysis, RNAseq, dPCR | Yes | CD4<sup>+</sup> T<sub>SCM</sub> | Figure 1, 2, 3, Supp 1, Supp 2 |
| 13 | 3 | Female | 70 | FACS analysis, RNAseq, dPCR | Yes | CD4<sup>+</sup> T<sub>SCM</sub> | Figure 1, 2, 3, Supp 1, Supp 2 |
| 86 | 4 | Female | 72 | FACS analysis, RNAseq, dPCR | Yes | CD4<sup>+</sup> T<sub>SCM</sub> | Figure 1, 2, 3, Supp 1, Supp 2 |
| 11 | 5 | Male | 28 | FACS analysis, dPCR, qPCR | Yes | CD4<sup>+</sup> total | Figure 3, Supp 4 |
| 10 | 6 | Female | 63 | FACS analysis, dPCR, CCL22-Trp treatment | Yes | CD4<sup>+</sup> total | Figure 3, Supp 6 |
| 75 | 7 | Male | 60 | FACS analysis, dPCR; qPCR | Yes | CD4<sup>+</sup> total | Figure 3, Supp 4 |
| 70 | 8 | Male | 18 | FACS analysis, dPCR, qPCR | Yes | CD4<sup>+</sup> total | Figure 3, Supp 4 |
| 58 | 9 | Female | 36 | FACS analysis, CCL22-Trp treatment | Yes | CD4<sup>+</sup> total | Supp6 |
| 64 | 10 | Female | 40 | FACS analysis | No | No | Figure 1, Supp 1 |
| 38 | 11 | Unknown | Unknown | FACS analysis | No | No | Figure 1, Supp 1 |
| 12 | 12 | Male | 25 | FACS analysis | No | No | Figure 1, Supp 1 |
| 47 | 13 | Unknown | Unknown | FACS analysis of CCL22 and IDO1 in subsets | No | No | Figure 4 |
| 54 | 14 | Unknown | Unknown | FACS analysis of CCL22 and IDO1 in subsets | No | No | Figure 4 |
| 83 | 15 | Unknown | Unknown | FACS analysis of CCL22 and IDO1 in subsets | No | No | Figure 4 |
| 89 | 16 | Female | 44 | CCL22-Trp treatment | No | No | Supp 6 |
| 12 | 17 | Male | 26 | FACS analysis | No | No | Figure 1 |

**Supplementary Table 2.**
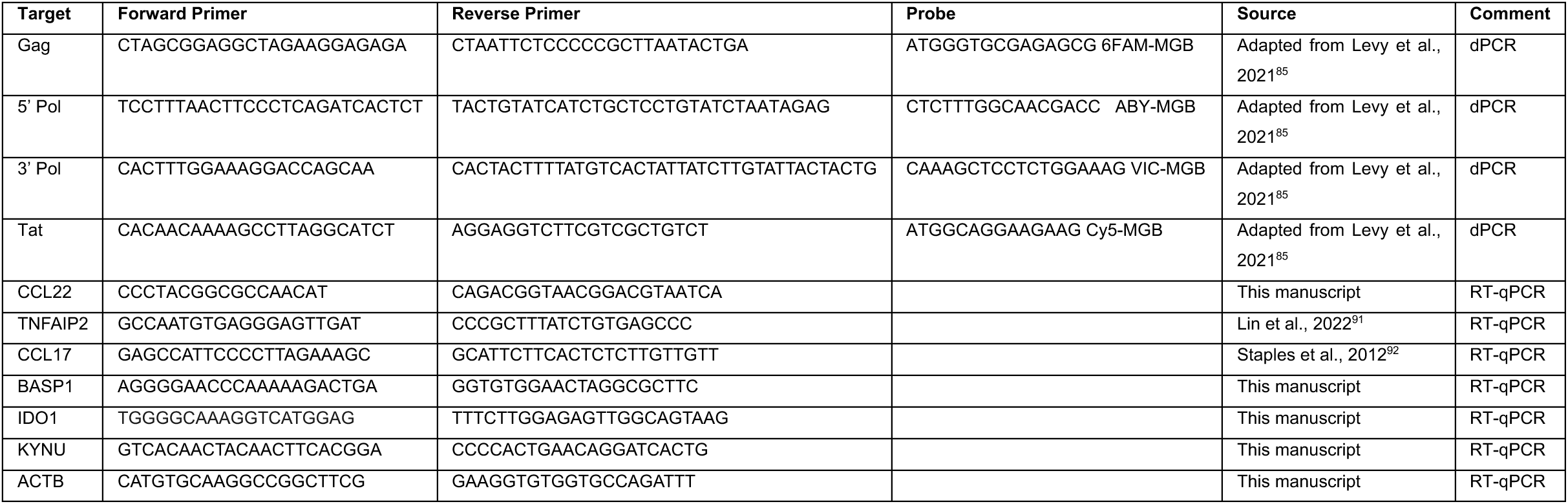
qPCR primer sequences. The table shows the sequences of the primers used for dPCR-PIA and RT-qPCR experiments

**Supplementary Figure 1.**
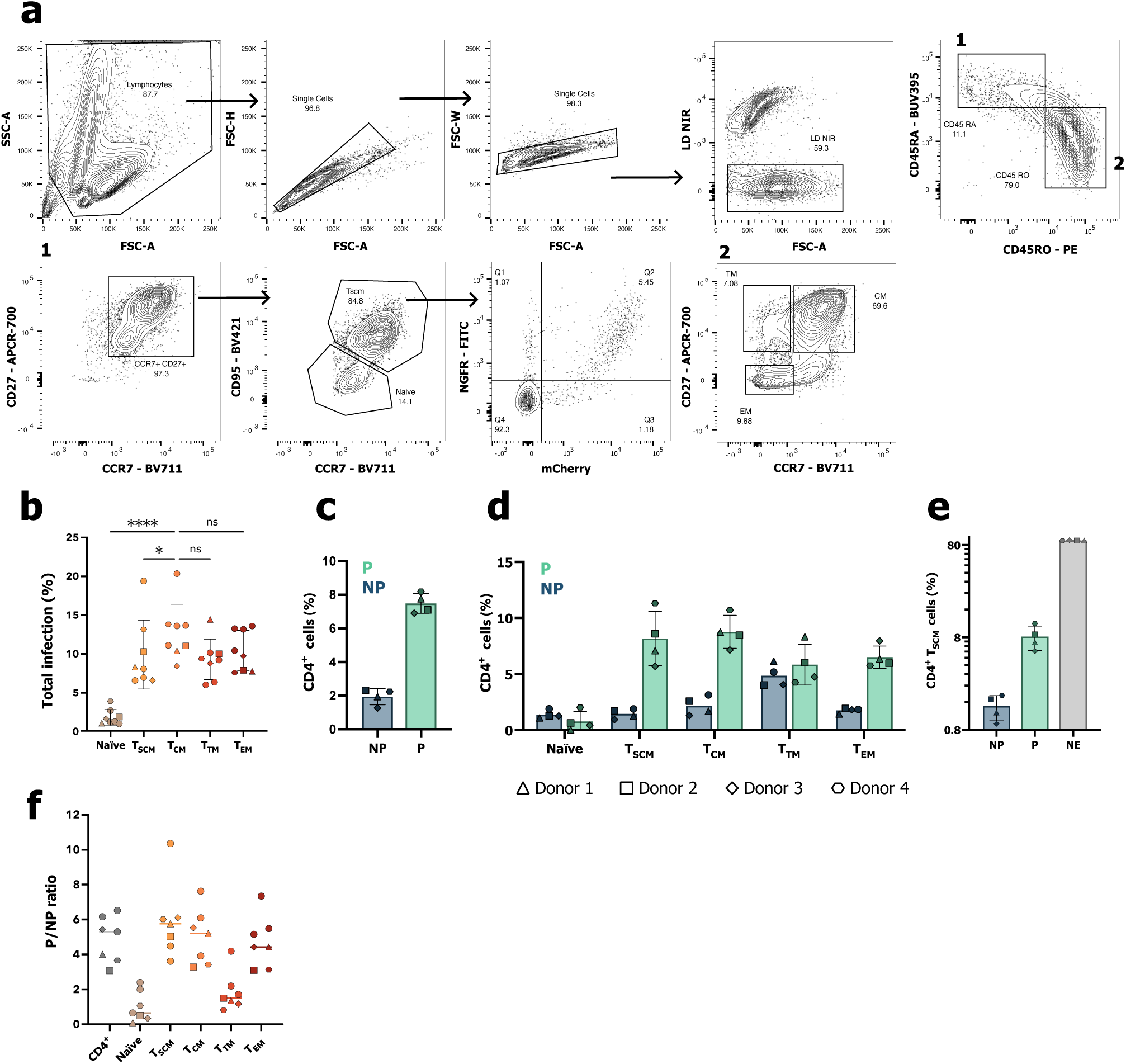
CD4^+^ T_SCM_ sorting strategy. (a) Representative gating strategy of purified peripheral blood CD4^+^ T cells infected with HIV pMorpheus-V5 reporter virus, at the time of sorting (five days post-infection). Total cultured CD4^+^ T cells were gated using physical parameters, then gated for Live Dead NIR and subsequently for CD45RA/CD45RO (“1” and “2”). CD45RA^+^ cells (“1”) were gated for CD27/CCR7. CD27^+^CCR7^+^ cells were gated for CD95/CCR7: CD95^-^CCR7^+^ cells were considered CD4^+^ naïve cells, CD95^+^CCR7^+^ cells were considered CD4^+^ T_SCM_. NP CD4^+^ T_SCM_ cells were NGFR^+^mCherry^-^(Q1), P cells were NGFR^+^mCherry^+^ (Q2), NE cells were NGFR^-^mCherry^-^ (Q4). CD45RO cells (“2”) were gated for CD27/CCR7: CD27^+^CCR7^+^ cells were considered T_CM_, CD27^+^CCR7^-^ cells were considered T_TM_, CD27^-^CCR7^-^ cells were considered T_EM_. (b) Dot plot representing total infection levels (NGFR^+^mCherry^-^+ NGFR^+^mCherry^+^) in CD4^+^ T naïve and memory subsets (n=8). Significance was calculated using one-way ANOVA (Dunnett’s multiple comparisons test, *, P < 0.05: **, P < 0.01, ***, P < 0.001, ****, P < 0.0001, ns, not significant). Each symbol represents a different healthy blood donor, with the four donors used for subsequent CD4^+^ T_SCM_ sorting highlighted with different symbols. Donors used for analysis but not sorted are represented by dots. (c) Percentages of non-productively (NP) and productively (P) infected cells in total CD4^+^ T cells and in (d) each CD4^+^ T cell subset considered, in the four donors sorted (each donor is represented by a different symbol, n=4). (e) Frequencies of CD4^+^ T_SCM_ NP, P and NE (negative-exposed) sorted populations in the four sorted donors (each donor is represented by a different symbol, n=4). (f) Dot plot representing the ratio between productively and non-productively infected cells in total CD4^+^ T cells and in the subsets considered (n=7). Primary CD4^+^ T cells from four different healthy donors (donors 1, 2, 3, 4) were stimulated with αCD3/CD28 antibodies for 72 hours before infection with HIV pMorpheus-V5 single-round reporter virus for five days before sorting. Data presented as the mean with SD.

**Supplementary Figure 2.**
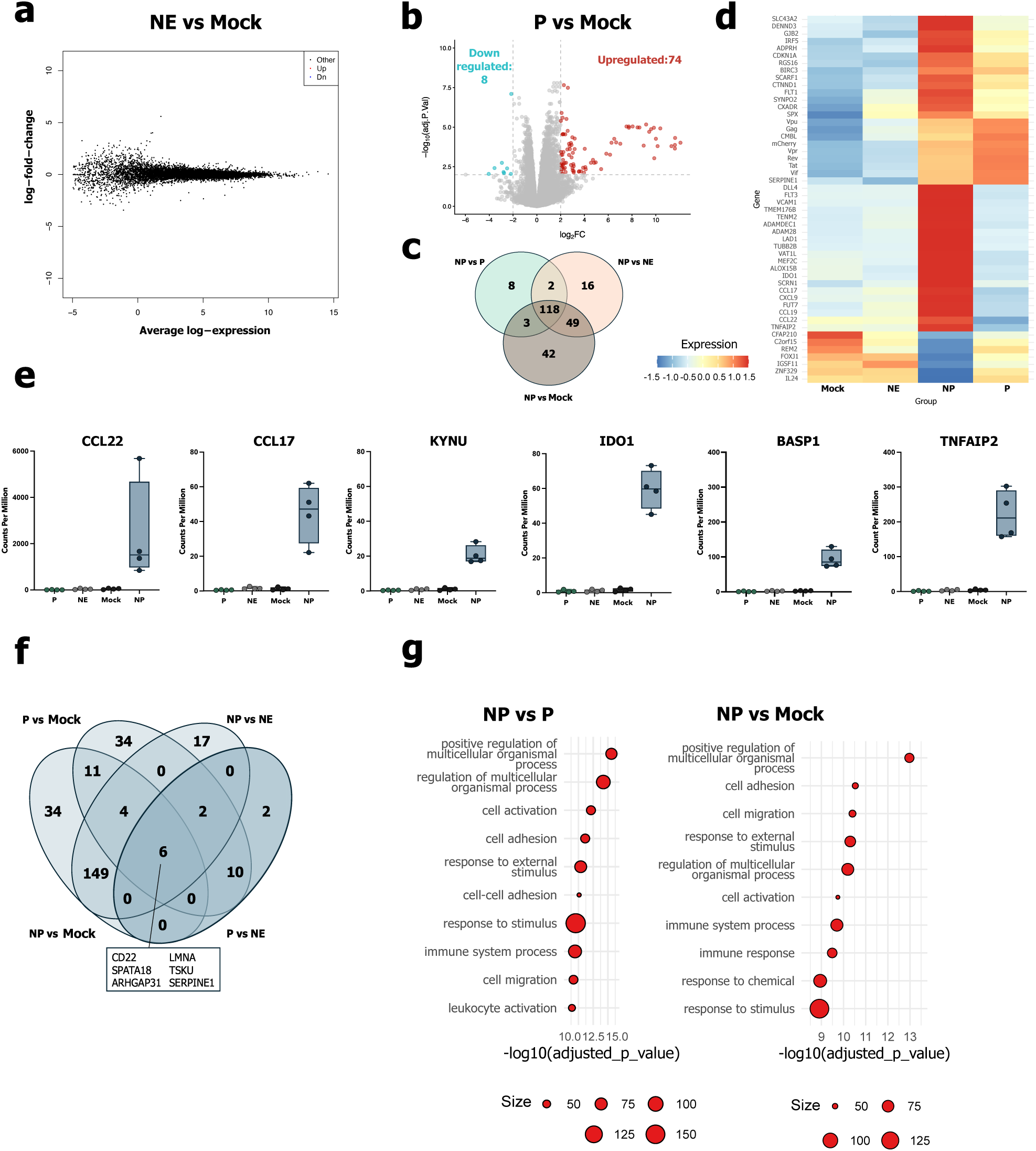
RNAseq DGE analysis of NP CD4^+^ T_SCM_ specific and CD4^+^ T_SCM_ virus-specific genes. (a) Mean-difference (MD) plot showing that no differentially expressed genes were found in CD4^+^ T_SCM_ NE compared to Mock (|log_2_FC| >= 2 and FDR <= 0.01). (b) Volcano plot representing the relative expression of all genes in the CD4^+^ T_SCM_ P vs mock comparison, with upregulated genes in red and downregulated genes in blue. (c) Venn diagram showing significantly upregulated genes in CD4^+^ T_SCM_ NP compared to P, NE and mock (log_2_FC >= 2 and FDR <= 0.01). (d) Heatmap of CD4^+^ T_SCM_ selected differentially expressed genes (Z-score is shown, row normalized considering only the displayed genes) across all sorted populations. (e) Boxplots representing transcript counts per million (CPM) detected by RNAseq in CD4^+^ T_SCM_ for genes encoding for *CCL22*, *CCL17*, *KYNU, IDO1*, *BASP1* and *TNFAIP2* in all sorted conditions (n=4). (f) Venn diagram of significantly upregulated genes in CD4^+^ T_SCM_ harboring an integrated pMorpheus-V5 provirus (either NP or P compared to NE and mock, log_2_FC >= 2 and FDR <= 0.01). The six shared significantly upregulated genes are highlighted in the box below the Venn diagram. (g) Gene Ontology (GO) of Biological Processes (BP) of significantly upregulated genes in NP CD4^+^ T_SCM_, compared to P (left) and mock (right).

**Supplementary Figure 3.**
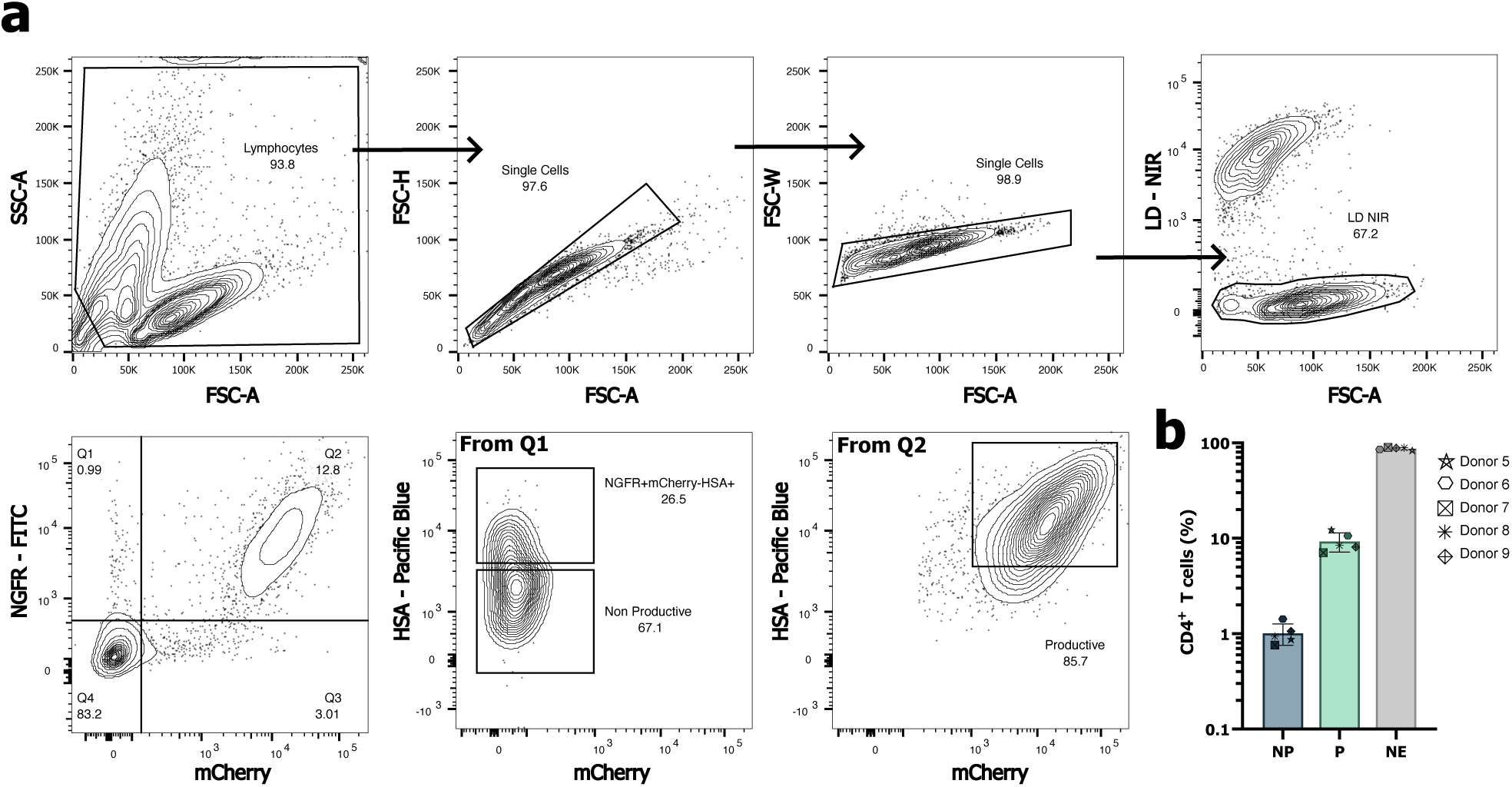
Total CD4^+^ T cell sorting strategy. (a) Representative gating strategy of purified peripheral blood CD4^+^ T cells infected with HIV pMorpheus-V5 reporter virus, at the time of sorting (five days post-infection). Total cultured CD4^+^ T cells were gated using physical parameters, then negative for Live Dead NIR, then gated for NGFR and mCherry. NGFR^+^mCherry^-^ cells were gated for HSA: NGFR^+^mCherry^-^HSA^-^ cells were considered CD4^+^ T cells non-productively infected with HIV pMorpheus-V5. NGFR^+^mCherry^+^ were gated for HSA: NGFR^+^mCherry^+^HSA^+^ were considered CD4^+^ T cells productively infected with HIV pMorpheus-V5. (b) Percentages of non-productive, productive infections and negative-exposed in total sorted CD4^+^ T cells, in the five sorted donors (donors 5-6-7-8-9, n=5). Each sorted donor is represented by a different symbol. Primary CD4^+^ T cells were stimulated with αCD3/CD28 antibodies for 72 hours before infection with HIV pMorpheus-V5 single-round reporter virus. Data presented as the mean with SD.

**Supplementary Figure 4.**
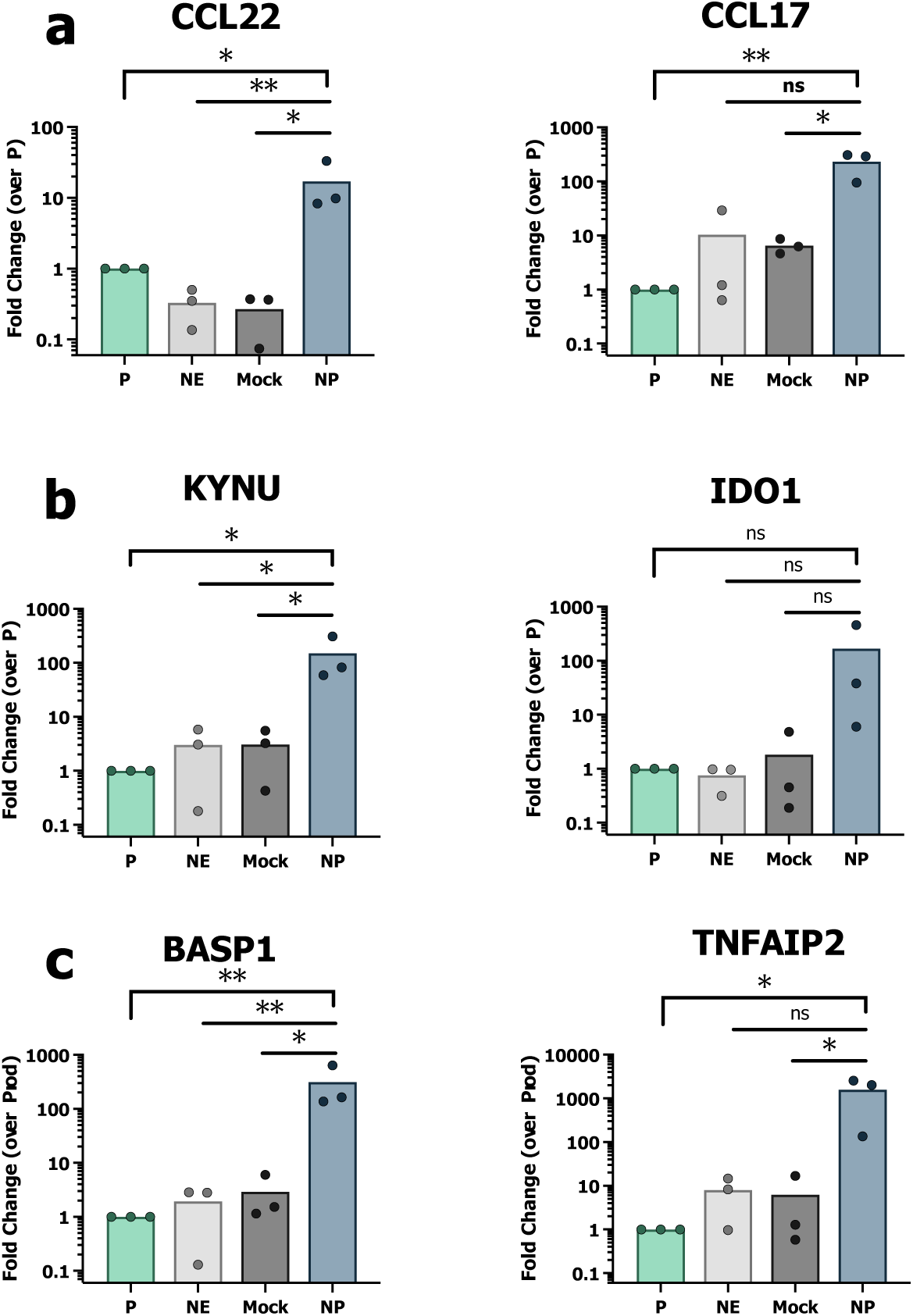
qPCR validation (in total sorted CD4^+^ T cells) of top hit genes upregulated in NP CD4^+^ T_SCM_. (a) Expression of *CCL22* and *CCL17,* (b) *KYNU* and *IDO1,* (c) *BASP1* and *TNFAIP2* mRNA in FACS-sorted total CD4^+^ T cells (Productively infected P, negative-exposed NE, mock unexposed mock, non-productively infected NP, compared to P) infected with HIV pMorpheus-V5 reporter virus (n=3). Significance was calculated on log-transformed fold-change values (normalized to P fold-change values, set as 1) using both one-way ANOVA with multiple comparisons (Dunnett’s multiple comparisons test, between NP, NE and mock) and one-sample t and Wilcoxon test (between single populations compared to P, only NP vs P t test significance is shown). *, P < 0.05: **, P < 0.01, ***, P < 0.001, ****, P < 0.0001, ns, not significant. Each dot represents a different donor.

**Supplementary Figure 5.**
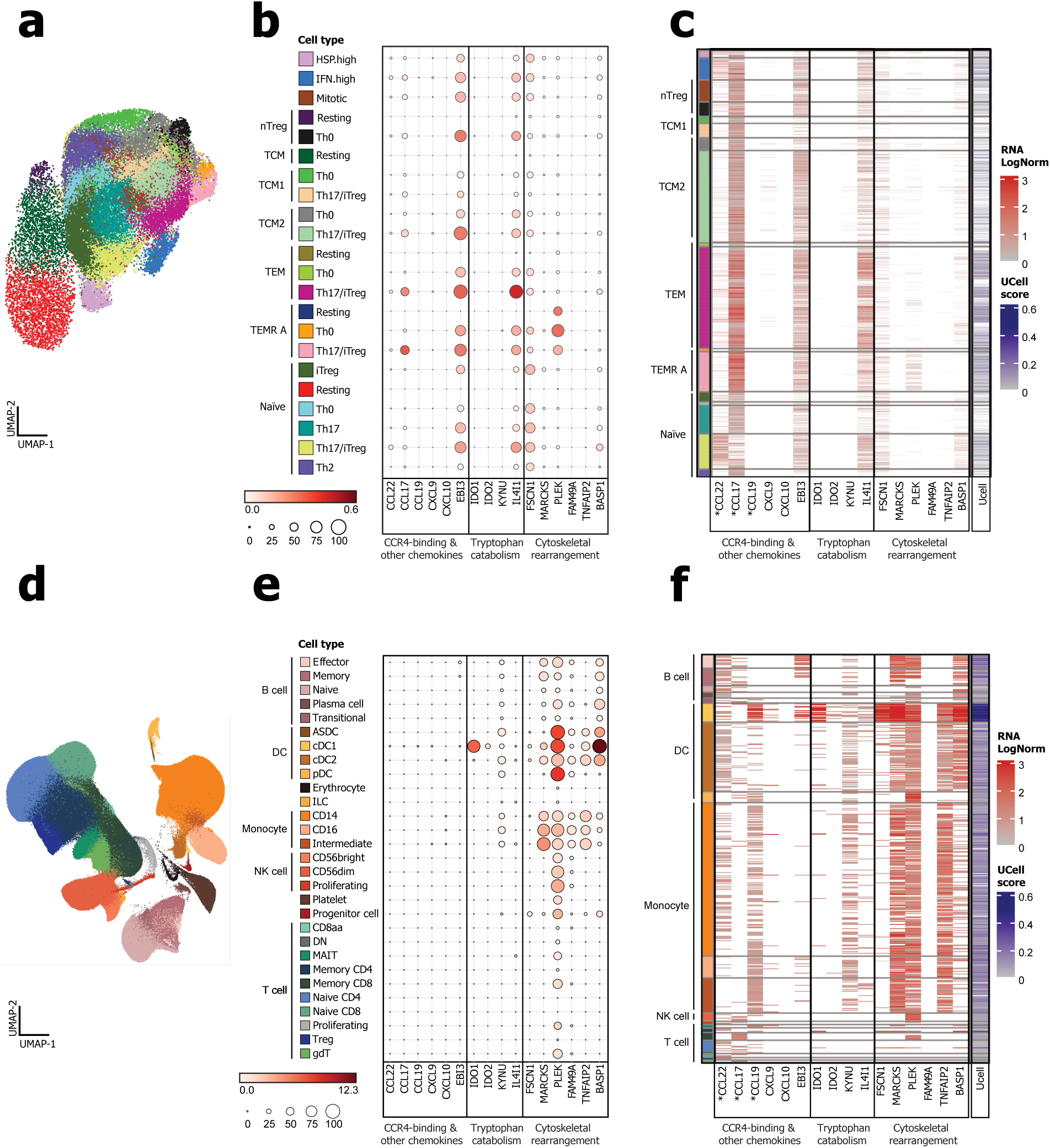
Gene expression and enrichment analysis of non-productive CD4^+^ T_SCM_ signature genes in human immune cell populations. (a) Uniform manifold approximation and projection (UMAP) of 43,112 naïve (TN) and memory (TM) CD4^+^ T cells from Cano-Gamez et al. (2020)^43^. Cell type populations in the UMAP are colored according to the legend: naïve, central (T_CM_) and effector (T_EM_) memory cells, effector memory cells re-expressing CD45RA (T_EMRA_), natural T regulatory (nTreg). (b) Dot plot showing the average gene expression per cell type and respective percentage of cells expressing each of 16 NP-upregulated genes, including chemokines (*CCL22, CCL17, CCL19, CXCL9, CXCL10, EBI3*), tryptophan catabolic enzymes (*IDO1, IDO2, KYNU, IL4I1*), and cytoskeletal regulators (*TNFAIP2, BASP1, FSCN1, MARCKS, PLEK, CYRIA*). (c) Heatmap displaying the expression levels of the 16 genes and corresponding UCell enrichment scores^90^ at the single-cell level in 2,756 CD4^+^ T cells (6.4%) in cells with detectable levels of *CCL22*, *CCL17* or *CCL19* (marked by *). (d) UMAP of 1,821,725 human immune cells within the Human Immune Health Atlas (T cells, B cells, monocytes, natural killer (NK) cells, and 12 other subsets including dendritic cells (DC) and hematopoietic precursors)^44^. Cell type populations in the UMAP are colored according to the legend. (e) Same as panel (b) but for the Human Immune Health Atlas^44^. (f) Same as panel (c) but for the 448 cells (0.025%) with detectable *CCL22*, *CCL17* or *CCL19* expression in the Human Immune Health Atlas (marked by *)^44^.

**Supplementary Figure 6.**
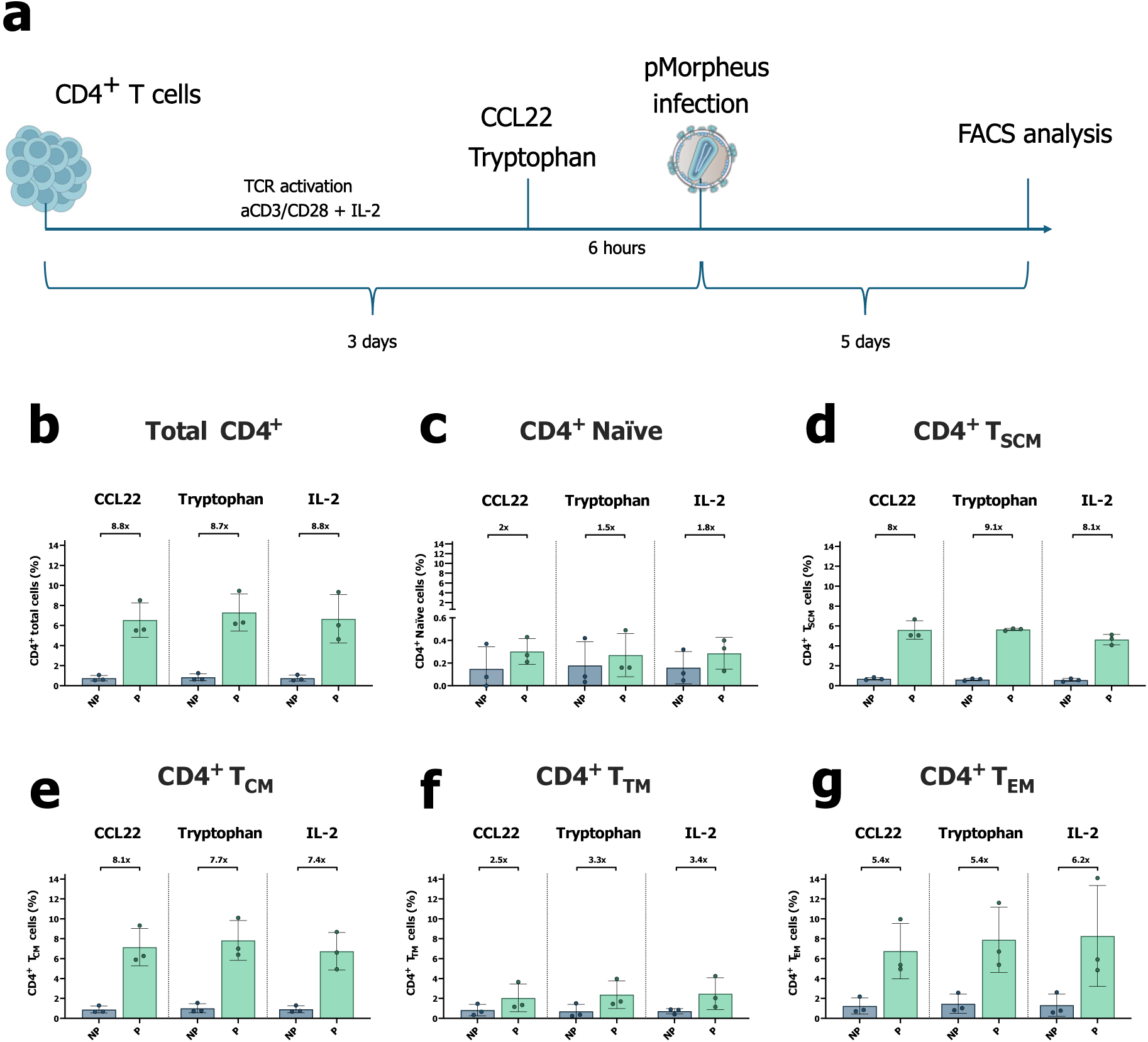
CCL22 or tryptophan treatment does not perturb latency establishment. (a) Experimental outline of CD4^+^ T cells treated with either CCL22, Tryptophan or IL-2 (media control). CD4^+^ T cells were TCR activated for three days in the presence of IL-2. Six hours before spinoculation, CCL22 or tryptophan was added to the cells. After spinoculation, cells were kept in either CCL22, tryptophan or control media until FACS staining and analysis. (b through g) Bar graphs showing the percentages of NP and P cells in the conditions treated with CCL22, tryptophan or IL-2 (media control), in total CD4^+^ T cells (b), naïve (c), T_SCM_ (d), T_CM_ (e), T_TM_ (f) and T_EM_ (g, n=3), with the ratio between P and NP highlighted on each condition (P/NP).

## Notes

### Competing Interest Statement

The authors have declared no competing interest.

